# Beta_2_ -Adrenergic Agonists in Treatment for Parkinsonism, with Implications for Neurodegenerative and Neoplastic Disorders

**DOI:** 10.1101/2024.01.12.575406

**Authors:** Mario A. Inchiosa

## Abstract

There is a long record of observations suggesting that β_2_-adrenergic agonists may have therapeutic value in Parkinson’s disease. Recent studies have focused on the possible role of β_2_-receptor agonists in suppressing the formation of α-synuclein protein, the component of Lewy bodies. Levalbuterol, the pure levo isomer of the β2 selective agonist, albuterol, has been found to possess significant anti-inflammatory activity, a property that may have the potential to suppress cytokine mediated degeneration of dopaminergic neurons and progression of Parkinsonism. All the β2 agonist and anti-inflammatory activities of albuterol reside in the levo isomer. The dextro isomer of albuterol substantially negates the efficacies of the levo form. Epinephrine, the prototypical β2 agonist and certain other adrenergic agents, when modeled in the Harvard/MIT Broad Institute genomic database, CLUE, demonstrated strong associations with the gene-expression signatures of drugs possessing glucocorticoid receptor agonist activity. Gene-expression signatures generated by the interaction of the adrenergic drugs of interest in 8 human tumor cell lines were compared with the entire CLUE database of more than 8,000 agents. The signatures were summarized for their consistency (connectivity) across all 8 cell lines and ranked for their relative degree of similarity to the agents in the database. Possible associations with anti-inflammatory activity of glucocorticoids prompted *in vivo* biological confirmation for levalbuterol and related agonists in the Jackson Laboratory human peripheral blood mononuclear cell (PBMC)-engrafted mouse. Levalbuterol inhibited the release of the eosinophil attractant chemokine, eotaxin-1 (specifically, CCL11), when the mice were challenged with mononuclear antibodies known to provoke cytokine release. Eotaxin is implicated in CNS and peripheral inflammatory disorders. Also, elaboration of the broad tumor-promoting angiogenic factor, VEGFa, and the pro-inflammatory cytokine, IL-13, from activated PBMCs were also inhibited by levalbuterol. These observations suggest possible translation to Parkinson’s disease, other neurodegenerative syndromes, and malignancies, by several mechanisms.

## 1. Introduction

Magistrelli and Comi [1] have reviewed the earlier history of reports of an association between treatment with β_2_-adrenergic receptor (β2AR) agonists and a therapeutic benefit in patients with Parkinson’s disease (PD). Three open-label studies were noted, dating from 1992, 1994 and 2003, with the use of salbutamol as an adjunct to levodopa in a total of 25 patients. In the earliest study (9 PD patients), there were significant increases in “on time” and shortening of response latency [2]. The 1994 study with 8 PD patients [3] showed statistically significant improvement in PD tests for manual dexterity and time to arise and walk 20 feet and return and sit again; the 2003 study with 8 PD patients also found improvement in manual dexterity and in the Unified Parkinson’s Disease Rating Scale (UPDRS) scores. [4]

### 1.1. α-Synuclein gene (SNCA) and PD

The observations in these earlier studies have advanced considerably with the recognition that β2AR ligands modulate the transcription of the α-synuclein gene (*SNCA*); and, the gene product, α-synuclein (α-syn), is implicated in the development of PD [1,5]. PD pathology is clinically correlated with the extent of excessive expression of *SNCA* and synthesis of α-syn, which accumulates intracellularly in Lewy bodies in the brains of PD patients.

In addition to PD, synucleinopathies include the neurodegenerative diseases Lewy bodies dementia and multiple system atrophy. All three demonstrate cellular accumulation of α-syn and loss of dopaminergic neurons [6].

Mittal et al. [5] screened 1126 FDA approved drugs and a variety of health supplements in cultured human neuroblastoma cells (SK-N-MC) for their ability to reduce transcription of *SNCA* mRNA in a four-stage gene expression assay. The highest efficacies were found in 3 β2AR agonists, metaproterenol, clenbuterol and salbutamol and, interestingly, in the drug, riluzole, which has been approved by the FDA for intervention in amyotrophic lateral sclerosis, where it was found to decrease dopaminergic degeneration in a PD model in the rat. All three β2AR agonists caused statistically significant relative reductions for *SNCA* mRNA abundance and α-syn protein abundance in SK-N-MC cells [5]. Of particular relevance to PD pathology and loss of dopaminergic neurons, these investigators also found that intraperitoneal clenbuterol treatment for 24 hours reduced expression of *SNCA* in the substantia nigra of C57BL/6J wild type mice, with decreases in nigral mRNA and α-syn protein. Additionally, higher cellular levels of α-syn produced mitochondrial dysfunction and production of superoxide anions and other reactive oxygen species (ROS); clenbuterol treatment in neuronal precursor cells (iPSC-derived) from a PD patient suppressed these changes [5].

### 1.2. PD causation from neuroinflammation

A number of studies have focused on the importance of inflammatory processes in neurodegenerative diseases, including PD. In this connection, there is considerable interest in the apparent role of β2AR signaling on immune cells in suppressing the inflammatory potential of microglia and macrophages, and in T cells and B cells. Signaling through this receptor can influence the inflammatory response of these cells [1,5,7–13].

The role of neuroinflammation in the initiation and progression of PD is a major focus of these present investigations. The functional role of the β2AR in biological responses has been further elucidated by the development of “short-acting” and “long-acting” relatively selective β2AR agonists that have little β1AR efficacy. These drugs have been used primarily for the treatment of bronchial asthma and chronic obstructive pulmonary disease (COPD). The nomenclature for these agents in the literature includes either the “United States Adopted Name” (USAN) or the “International Nonproprietary Name” (INN), usually depending upon the country of origin of the research or the date of the publication. The designated names of the principal drugs that were investigated are as follows: Albuterol sulfate is the USAN for a short acting bronchodilator that has an INN of salbutamol sulfate. Albuterol is a racemic mixture of equal quantities of the S and R isomers; all of the β2-adrenergic activity resides in the R isomer. Some proprietary names for albuterol sulfate are Ventolin, Proventil and ProAir. The pure R-isomer has the USAN of levalbuterol HCl (a proprietary formulation as levalbuterol tartrate is named Xopenex); its INN is levosalbutamol (it is also referred to as R-salbutamol). Arformoterol is the USAN for the pure R isomer of a long-acting β2AR agonist; the racemic mixture of the drug is named formoterol. The bronchodilator drug theophylline was also studied; it does not have β2AR agonist activity but inhibits the metabolism of cyclic AMP (cAMP) by phosphodiesterase enzymes thereby preserving intracellular levels; cAMP is the initial signaling messenger that results from activation of both β1- and β2-ARs.

It is important to note the differences in the expected anti-inflammatory activity of β2AR agonists regarding whether they are racemic mixtures, such as albuterol, or the pure R-isomer, such as levalbuterol. In addition to the fact that the R-isomer has all the bronchodilator activity of albuterol, it also has all the anti-inflammatory activity. The S-isomer of albuterol has pro-inflammatory activity and partially, completely, or overwhelmingly negates the anti-inflammatory activity of the R-isomer. Mazzoni et al. [14] demonstrated a hyperresponsiveness to histamine in guinea pig airways following pretreatment with S-albuterol. This is apparently related to the capacity of that isomer to increase smooth muscle proliferation, as observed in human bronchial cell cultures [15]; levalbuterol decreased cell proliferation compared with the control cultures enriched with 5% fetal bovine serum. Baramki et al. [16], Chorley et al. [17] and Wang et al. [7] have all confirmed the anti-inflammatory effects of levalbuterol (R-albuterol) and the pro-inflammatory and compromising effects of S-albuterol on those of levalbuterol.

### 1.3. Epidemiological associations between beta2-adrenergic effects and PD

Several large epidemiological studies have revealed the finding that the clinical use of β2AR agonists (primarily for patients with asthma) had a decreased incidence of PD compared to the general population and that the use of a β2AR antagonist (primarily for the treatment of hypertension) had an increased incidence. Mittal et al. [5] studied the Norwegian Prescription Database over an 11-year period for the entire population alive on January 1, 2004 (n = 4.6 million). Patients treated with salbutamol (albuterol) showed a decreased PD risk; the rate ratio (incidence with salbutamol/incidence without exposure to salbutamol) was 0.66 (95% confidence interval, CI of 0.58 to 0.76). Patients treated with propranolol, a β1-,β2AR antagonist had a marked increased rate ratio of PD to 2.20 (95% CI of 1.62 to 3.00).

Using health records from the major Israeli health provider, Gronich et al. [18] studied medical records of 1,762,164 individuals who did not have a PD diagnosis on January 1, 2004 and followed their health histories until June 30, 2017. Each participant who was treated with salbutamol was compared with 10 cohorts as controls and showed a decreased rate ratio for development of PD of 0.89 (0.82-0.96); p = 0.004. Participants treated with propranolol showed an increased PD rate ratio of 2.60 (2.40-2.81); p <0.001. A special feature of this study [18] was that it included a number of participants that were treated with a group of β1 selective antagonists instead of the β1-, β2AR antagonist, propranolol. A total of 3,032 participants received either metoprolol, atenolol or bisoprolol. Since these drugs leave β2ARs largely available, it would be expected that they would be chronically stimulated by endogenous epinephrine that was released during daily activities and exercise. These participants showed no change in PD rate ratio as compared with their cohort controls; 1.00 (0.95-1.05); p = 0.94. All the above epidemiological studies consistently demonstrated that drugs that stimulated the β2AR receptor, or interventions that left it available for activation, decreased the incidence of PD.

Chen et al. [19] conducted a meta analysis on the association between β-adrenoceptor drugs and Parkinson’s disease and distilled 640 studies down to 8 that were appropriate for meta-analysis. Salbutamol yielded a decreased rate ratio for PD of 0.888 (0.822-0.960); p = 0.003. When the studies that used one or more of the β2AR agonists were analyzed together, the rate ratio for PD was still lowered: 0.840 (0.714-0.987); p = 0.035. This included the β2 selective agonists salbutamol, formoterol, salmeterol, and terbutaline. It must be noted that all these drugs are the racemic mixtures; in the studies noted above, only the levo (R) isomers have been shown to have anti-inflammatory activity while the S-isomer portion is actually proinflammatory and compromises the effects of the R-isomer. Thus, the decreased incidence of PD in participants that received β2AR agonists may have been greater if the single R-isomer agents had been studied.

### 1.4. Life-style factors that may influence the incidence or progression of PD

There is considerable evidence that certain lifestyle practices delay the initiation of PD and the rate of its progression. These include physical exercise, nicotine exposure from smoking, and intake of caffeine from consumption of coffee or tea. In a potential connection with the present study, these life-style interventions are associated with an increased outflow of the sympathetic nervous system and elevations in circulating levels of epinephrine, with its prominent β2AR agonist activity.

Exercise programs are a major therapeutic recommendation on the websites of the Parkinson’s Foundation and the Michael J. Fox Foundation to forestall the progression of disease and to ameliorate the symptoms of PD. Moore et al. [20] reviewed extensive literature showing the beneficial effects of physical exercise in human and animal models of PD. They also presented the projected design of a Phase II trial, “Study in Parkinson Disease of Exercise,” (SPARX). It was intended to serve as a pilot study to determine whether a larger study on graded intensity of exercise was warranted. It included patients who were at an early stage of PD and were not on dopaminergic therapy or, if so, for only a short period. There were 3 groups: High intensity treadmill exercise for 30 min, 4 times/week at 80-85% of their maximum heart rate; moderate intensity exercise for 30 min, 4 times/week at 60-65% of maximum heart rate; and usual care for a wait-list group who were eligible later for inclusion in the study. The ability to identify treatment effects was complicated by the fact that the usual care group were permitted to continue their current exercise programs since, as noted above, the value of exercise in PD was already appreciated and it would not have been ethical to prevent the control group from possible benefits of their existing programs. The groups were to be maintained and monitored for a period of 6 months.

Schenkman et al. [21] reported the results of the Phase II pilot trial that included a total of 128 participants. There were improvements in the Movement Disorder Society revised Unified Parkinson Disease Rating Scale (MDS-UPDRS) in both treadmill groups, but statistical significance was only reached at the high intensity level. Recruitment is now ongoing for the SPARX3 trial that will be conducted at 29 sites in the United States and Canada (Study No. NCT04284436; sparx3pd.com) with 370 participants. This study will have considerably more statistical power since there will only be two groups divided at the two intensity levels; the changes in MDS-UPDRS scores will be evaluated for each participant between their initial values and at the end of the study period. The SPARX3 trial has been delayed by the COVID-19 pandemic and completion of recruitment is now estimated to be July, 2025, with the results available approximately 18 months after close of the study.

The studies of Kjaer et al. [22] on epinephrine release with exercise are of relevance in relation to the evolving focus on the intensity of exercise as therapy in PD. Their findings in a series of studies showed a direct relationship between exercise intensity in healthy adults and epinephrine arterial plasma concentrations. Four levels of exercise intensity were studied based on the individual subjects’ maximal oxygen uptake (VO_2_MAX). Resting epinephrine concentration averaged 0.5 nmol/l. Plasma epinephrine concentration after 60 min at 40-56% VO_2_MAX averaged 1.6 nmol/l; after 25 min at 60-70% VO_2_MAX, 2.2 nmol/l; after 25 min at 82-89% VO_2_MAX, 4.5 nmol/l; and after 4 min at 100-110% VO_2_MAX, 8.7 nmol/l.

Tobacco smoking has a long and established negative association with the onset and progression of PD [23–27]. Meta-analysis by Hernan et al. [27] that included 44 case-controlled studies and 4 cohort studies showed a relative risk of PD between current smokers and individuals that had never smoked of 0.39 (95% confidence interval of 0.32-0.47). The pharmacological effects of nicotine inhalation on the sympathetic nervous system would appear to be a convincing explanation for this protective effect. In addition to stimulating outflow of the sympathetic nervous system from the brain, nicotine activates nicotinic receptors in sympathetic ganglia peripherally to release norepinephrine from sympathetic nerve endings and also generates release of epinephrine by stimulation of nicotine receptors in the adrenal medulla.

Cryer et al. [28] studied the effects on catecholamine plasma concentrations in 10 male subjects in conjunction with two trials of smoking 5.5 cm of a cigarette in a 10 min period. The men were chronic smokers but had fasted overnight and had not smoked until the time of the experiments. Plasma concentrations of norepinephrine and epinephrine rose rapidly during the smoking period to maxima at about the 10-min time point. The average epinephrine concentration before smoking of 44 +/− 4 (mean; S.E.) pg/ml rose to 113 +/− 27 pg/ml (p < 0.05) at 10 min; it remained noticeably above the basal level for the next 20 min until measurements were halted. Grassi et al. [29,30] also studied the acute effects of cigarette smoking on epinephrine plasma levels. Their protocol required 48 hours of abstinence from smoking and completion of the cigarette in a 5 min period. Although the smoking period was half that in the previous study [28], epinephrine concentrations increased from 25 +/− 8.6 (mean; S.E.) pg/ml before smoking to 47 +/− 11.5 pg/ml (p <0.01) at the end of the period.

The relationship of caffeine consumption to a decreased incidence of PD has a long and consistent documentation [27,31–36]. Tan et al. [32], in a completely Chinese population, found that 10 years of coffee consumption at 3 cups/day was associated with a decreased odds ratio for PD of 0.78 (95% CI 0.66-0.93) p = 0.006, and for 10 years of tea drinking at 3 cups/day odds ratio of 0.72 (95% CI 0.56-0.94) p = 0.014). Ren and Chen [36] have reviewed the several large prospective and meta-analyses that show strong evidence of the negative association between caffeine consumption and PD. Notably, a meta-analysis by Qi and Li [35] that involved a total of 901,764 participants from 13 studies showed a maximum benefit in PD incidence reduction at approximately 3 cups of coffee/day: relative risk of 0.72 (95% CI 0.65-0.81). This same review [36] presented evidence from animal studies that caffeine may reduce neuroinflammation and its attendant loss of dopaminergic neurons. Also, in α-syn animal models of PD, caffeine administration was associated with a decreased aggregation of α-syn, the component of Lewy bodies.

Finally, consistent with the above observations with physical exercise and cigarette smoking, caffeine consumption results in increases in plasma epinephrine. In a precisely controlled clinical study, Robertson et al. [37] demonstrated that 250 mg of caffeine in non-coffee drinkers resulted in a significant increase in plasma epinephrine concentration one hour after consumption from 36 +/− 5 pg/ml (mean, S.E.) for a control beverage to 89 +/− pg/ml with caffeine; p <0.001. Plasma epinephrine concentrations were still statistically different from controls in the final measurements 3 hrs. after caffeine consumption.

Benowitz et al. [38] observed similar findings in caffeine consumption of 2 and 4 mg/kg with a cross-over design in coffee drinkers who had abstained for 3 days before the study. The results, expressed as maximum changes in epinephrine concentrations were 35 +/− 19 (mean, S.E.) pg/ml for the 2 mg/kg dose and 63 +/−18 pg/ml for 4 mg/kg. Both doses were statistically different from controls (p <0.05) and the 4 mg/kg dose was statistically greater than the 2 mg/kg dose (p <0.05). Measurements were made over a 180 min period and the areas under the time x plasma concentrations (AUCs) were also statistically and markedly different from the controls, with the 4 mg/kg dose again statistically different from the 2 mg/kg dose (differences of p <0.05).

### 1.5. Genomic modeling of the potential anti-inflammatory effects of epinephrine

The evidence outlined above on the possible role of β2AR agonist activity on: 1, Reducing the expression of *SNCA* mRNA and its protein product α syn; 2, The reduction in neuroinflammation and loss of dopaminergic neurons; 3, Epidemiological associations with decreased incidence of PD in their routine clinical use; and 4, Life-style factors such as physical exercise, cigarette smoking and caffeine consumption that are negatively associated with PD incidence, and which all result in increased plasma epinephrine concentration, prompted an attempt to explore a possible link between the gene-expression characteristics of epinephrine and an anti-inflammatory potential.

Epinephrine was modeled in the Harvard/MIT Broad Institute genomic database, CLUE [39]. It was found to have a remarkably coherent similarity in gene-expression activity to the human glucocorticoid, cortisol (hydrocortisone), which has profound anti-inflammatory activity. Of particular interest, this property was not shared with norepinephrine, which is known to have minimal β2AR agonist activity. These observations led to the investigation of the potential anti-inflammatory effects of a series of β2AR agonists in the Jackson Laboratory human peripheral blood mononuclear cell (PBMC)-engrafted mouse.

### 1.6. Anti-inflammatory studies in the Cytokine Release Syndrome (CRS) assay

The CRS assay of Jackson Laboratories in humanized mice was utilized to evaluate the anti-inflammatory potential of several of the β2AR agonists and related compounds noted above, primarily to evaluate their capacity to protect against and modulate neuroinflammation. This assay employs immunodeficient mice that have been engrafted with human peripheral blood mononuclear cell cultures (PBMCs) that are then challenged with several monoclonal antibodies (mAbs) that are known to potently induce clinically injurious cytokine release [40,41]. The mice can be pre- and chronically treated with agents of interest for anti-inflammatory effects in conjunction with the mAb challenges. The Jackson Laboratories, Bar Harbor, Maine website provides additional details of the CRS assay.

The direct β2AR agonists, albuterol, levalbuterol, and arformoterol, and the indirect agonist theophylline, all noted above to increase cellular levels of cAMP, were studied in the mouse CRS assays. Phenoxybenzamine, a noncompetitive antagonist of alpha2-adrenergic agonist activity that results in increased cellular levels of cAMP [53], was also assayed. All the compounds except albuterol, the racemic mixture of S and R albuterol, showed anti-inflammatory potential in the CRS assay.

The possible anti-inflammatory activity of beta2-adrenergic agonists, as suggested by their gene-expression similarity to that of glucocorticoid receptor agonists, became a principal aim of these investigations. This hypothesis was tentatively confirmed by the inhibitory effects of beta2-adrenergic drugs on the release of several cytokines and chemokines in the activated human PBMC-engrafted mouse.

## 2. Materials and Methods

### 2.1. Analyses of gene-expression signatures of β2-adrenergic agonists and related compounds in the Harvard/MIT Broad Institute genomic database, CLUE

Both the earlier online Broad Institute CMap genomic dataset and the new cloud-based CLUE platform are based on the gene-expression signatures that result from the perturbation of actively proliferating cells that are malignant. The earlier online platform showed predictions of antitumor and histone deacetylase activity of phenoxybenzamine [56]. The strength of these predictions was extended when phenoxybenzamine was profiled on the cloud-based platform [57]. Both references provide additional details of the methodology.

The database and associated software are accessible at https://clue.io. The drugs, chemical agents and genetic inputs are referred to as, “perturbagens” [39, 42]. The present analyses focused on gene-expression signatures located in the “TOUCHSTONE” dataset (accessed in the “Tools” menu) that was selected because it represents a set of thoroughly annotated small molecule perturbagens that would be expected to be relevant to comparisons with the possible effects of β2-adrenergic agonists; see: https://clue.io/connectopedia/tag/TOUCHSTONE. The Touchstone dataset contains a total of approximately 8,400 perturbagens that have produced gene signatures that were generated from testing on a panel of 9 proliferating malignant cell lines; those cell lines are detailed in Table 1.

**Table 1.** Cell Lines Profiled in the TOUCHSTONE Database.

| Cell Line | Description |
| --- | --- |
| A375 | Human malignant melanoma |
| A549 | Human non-small cell carcinoma |
| HA1E | Human kidney epithelial immortalized |
| HCC515 | Human non-small cell lung adenocarcinoma |
| HEPG2 | Human hepatocellular carcinoma cell line |
| MCF7 | Human breast adenocarcinoma |
| PC3 | Human prostate adenocarcinoma |
| VCAP | Human metastatic prostate cancer |
| HT29 | Human colorectal adenocarcinoma |

In almost all instances, the perturbagens were tested at a concentration of 10 µM, allowing for the comparison of the strength of the connectivity of gene-expression between agents at equimolar concentrations.

It should be noted that the dataset in the earlier online platform of CMap was based almost entirely on perturbations in only 3 tumor cell lines, MCF7 (human breast adenocarcinoma), PC3 (human prostate adenocarcinoma) and HL60 (human promyeloblast); the first two cell lines are among the 9 in the current CLUE platform and consistency among the gene-expression signatures for the different cell lines is strengthened by the inclusion of additional cell lines.

The scoring value obtained in the gene expression analyses is termed “tau;” it ranges from 100 to −100 and is a measure of the connectivity between the gene expression-signature of the perturbagen of interest and those of the other 8400 perturbagens in the database. A positive tau indicates a relative similarity between two perturbagens or group of perturbagens, while a negative score indicates relative opposing gene signatures. Thus, for example, a tau score of 95 indicates that only 5% of the signatures in the database had connectivity that was higher than that of the perturbagen being profiled; see: https://clue.io/connectopedia/connectivity_scores. Positive scores above 90 are generally considered worthy of consideration as representing possibly important similarities in gene expression. The gene-expression data outputs from the analyses places emphasis on this range.

The Touchstone database returns two data formats in response to a query about the similarities or differences between a perturbagen of interest and the other perturbagens in the database. The compound being searched against all other entries in the Touchstone database is termed the “INDEX.” A “heatmap” format presents the connectivity score between the query perturbagen and the reference perturbagens in the database for each of the 9 cells lines that have been studied. (Seven to 9 cell lines may have been chosen for a particular INDEX analysis.) The second format is the “detailed list” output. In our experience, this output provided the most convenient access to information that relates to the primary focus of the present studies. The detailed list format also includes the protein targets of the individual perturbagens. Both formats provide comparison of the gene signatures of the INDEX compound with members of the four perturbagen classes in the Touchstone database. Those classes are identified as: Chemical compound/pharmacologic agent (CP); Gene knock-down (KD); Gene over-expression (OE); and perturbagen class (PCL). In the heat map format, INDEX signatures are compared with the entire Touchstone database of perturbagens (approximately 8400). Since our primary interest was to search for drugs and chemicals that had recognized mechanisms of action that might strengthen previous findings and support hypotheses for the repurposing of drugs therapeutically, we focused our searches in the detailed list format primarily to the CP agents.

The heatmap output that is presented gives the calculated *median* tau score (connectivity) for the 9 cell lines and a “summarization” measure. For all other results, the connectivity scores that are presented are the summarization measure. This measure is calculated across the 9 cell lines. It is valuable since it provides a measure of the consistency of perturbations from one cell line to another. The algorithms for the calculation of tau and the summarization measure, and for the associated formulas, are presented at https://clue.io/connectopedia/cmap_algorithms. Gene-expression connectivity scores are also associated with a “rank” measure for that score; the rank score has an almost perfect non-linear inverse correlation with the gene expression similarity/connectivity score. These measures are detailed further in S1 Figure.

The results represent the status of the CLUE platform when accessed primarily between May, 2020 and July, 2023; the database is dynamic and changes slightly if new perturbagens are added.

### 2.2. CRS assay methodology

Female NSG-SGM3 mice [*NOD.Cg-Prkdc**^scid^** Il2rg^tmlW**jl**^* Tg(CMV-IL3,CSF2,KITLG) lEav/MloySzJ;stock #013062] were used for the study. The mice were ear notched for identification and housed in individually ventilated polysulfonate cages with HEPA filtered air at a density of up to 5 mice per cage. Cages were changed every two weeks. The animal room was lit entirely with artificial fluorescent lighting, with a controlled 12 h light/dark cycle (6 am to 6 pm light). The normal temperature and relative humidity ranges in the animal rooms were 20-26°C and 30-70%, respectively. The animal rooms are set to have up to 15 air exchanges per hour. A 1% sucrose solution of filtered tap water, acidified to a pH of 2.5 to 3.0, and standard lab chow were provided ad libitum.

#### Study Design

1. Mice were irradiated and injected with PBMCs from a single donor on Day 0.

2. Animals were monitored for body weight and clinical observations once daily.

3. Animals were euthanized before study end if they showed >20% body weight loss or a body condition score of ≥2.

4. Treatment compounds were delivered in drinking water starting on Day 4 for the duration of study.

5. Mice were dosed on Day 6 with monoclonal antibodies OKT3 or anti-CD28, to induce cytokine release [41], or with PBS as control.

6. Mice were bled via retro-orbital (RO) method at 6 h post dose on Day 6. Blood was processed and measured for human cytokines and chemokines using MILLIPEX Human Cytokine/Chemokine/Growth Panel A Magnetic Bead Panel, Cat# HCYTOMAG-60K.

7. Bodyweight and clinical CRS score were assessed daily starting Day 6. Mice were euthanized by CO2 asphyxiation at 72 h post dose Day 9.

The experimental design of the study is summarized by Study Day (SD) as follows:

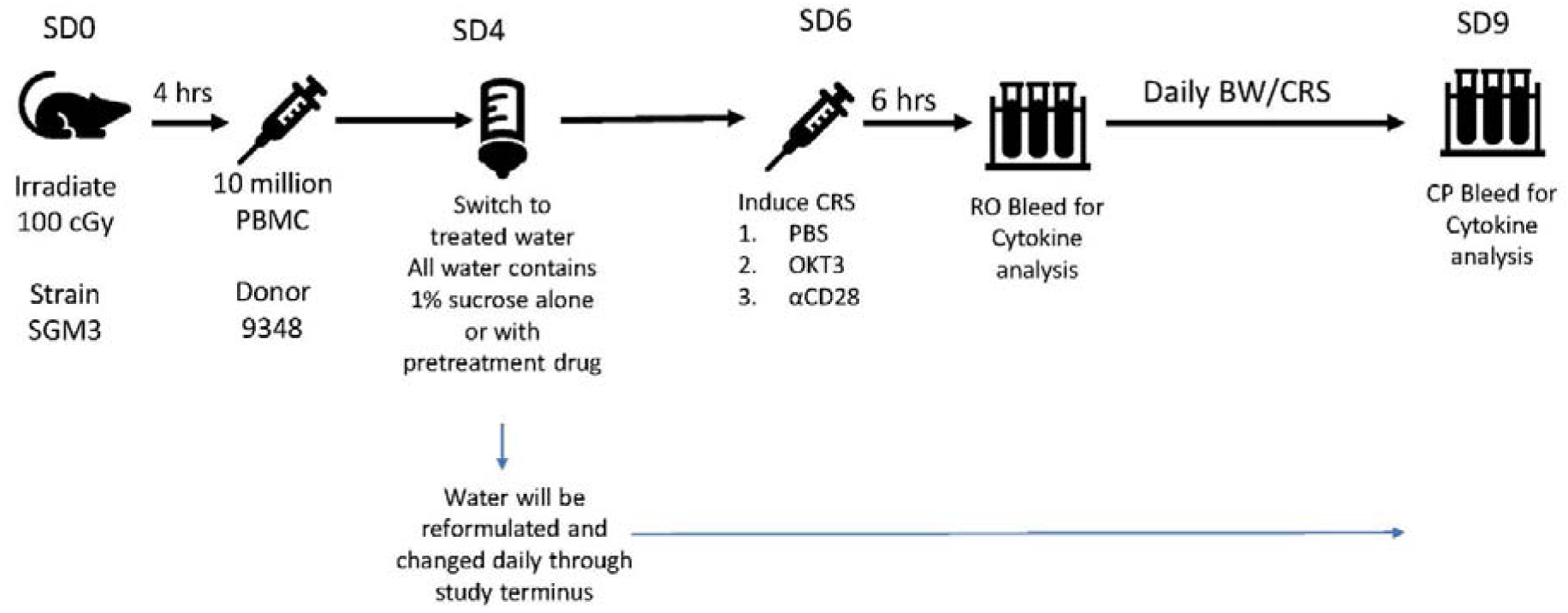

The clinical condition of the mice was rank-scored as follows:

<u>Clinical CRS score:</u>

0: normal activity

1: normal activity, piloerection, tiptoe gait

2: hunched, reduced activity but still mobile

3: hypomobile but mobile when prompted

4: moribund (non-responsive to touch)

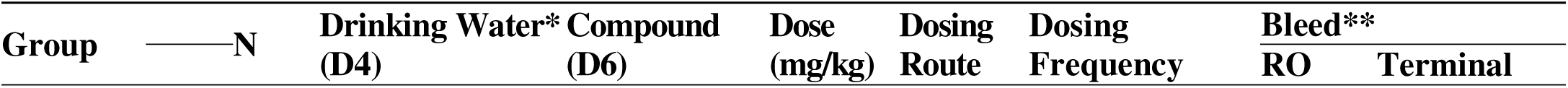
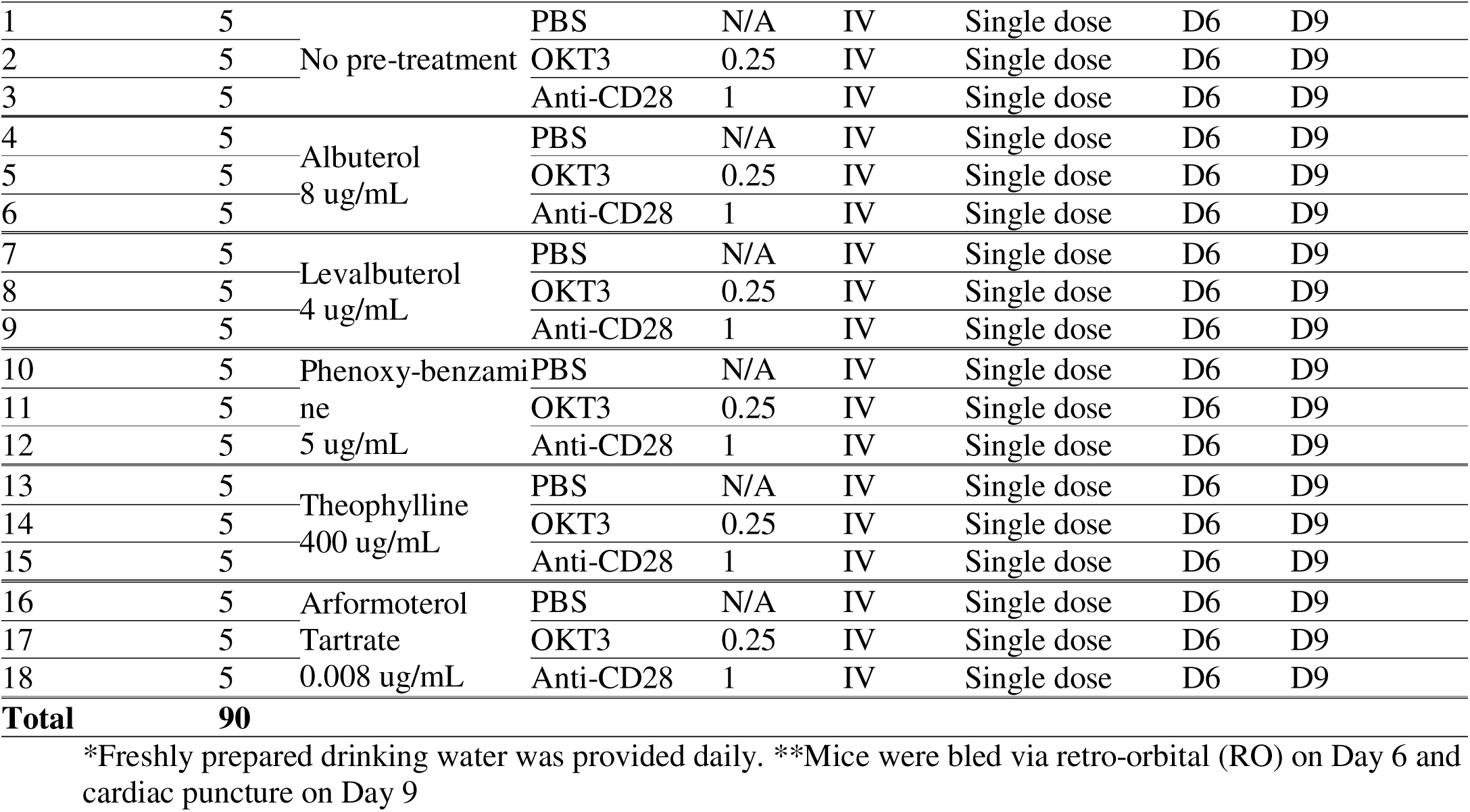

#### Dosing schedule and interventions

Levalbuterol Hydrochloride, Albuterol sulfate, and Phenoxybenzamine Hydrochloride were purchased as reference standards from the United States Pharmacopeia. Arformoterol Tartrate and Theophylline (anhydrous powder) were purchased from Sigma-Aldrich Chemical Company.

### 2.3. Statistics

Group sizes of these valued experimental animals were kept as small as possible and were consistent with the spirit of FDA Amendment 2.0. The cytokine and chemokine data from these small groups largely failed normality distributions. Thus, all analyses were conducted with the Kruskal-Wallis multiple comparative approach for nonparametric data followed by post-hoc adjustment of p values with the Dunnett’s test. Each drug treatment effect was only compared with its sucrose control. A p value of 0.05 (*) or less was considered statistically significant; ** p <0.01; ***p <0.001; ****p <0.0001.

## 3. Results and Discussion

### 3.1. Analyses of gene expression signatures of β2-adrenergic agonists and related compounds in the Harvard/MIT Broad Institute genomic database, CLUE

In preliminary analyses, salbutamol (albuterol), the racemic mixture of the drug that was prominent in the Norwegian and Israeli epidemiological studies in relation to PD frequency [5,18] was profiled in CLUE. As might be expected, salbutamol showed high connectivity with other beta-adrenergic receptor agonists; in this CMap class profiled in CLUE, 8 had gene-expression connectivity scores that were higher than 90.0 in relation to salbutamol (Table 2).

**Table 2.** Gene-expression connectivity scores of adrenergic receptor agonists in relation to salbutamol.

| Rank | Score | Name | Description | Target |
| --- | --- | --- | --- | --- |
| 3 | 99.93 | cimaterol | Adrenergic receptor agonist | ADRB3, ADRB1, ADRB2 |
| 27 | 99.37 | procaterol | Adrenergic receptor agonist | ADRB2 |
| 118 | 97.54 | dobutamine | Adrenergic receptor agonist | ADRB1, ADRB2, ADRA1A 3 |
| 183 | 96.72 | orciprenaline | Adrenergic receptor agonist | ADRB2 |
| 255 | 95.52 | buphenine | Adrenergic receptor agonist | ADRB2 |
| 288 | 95.21 | fenoterol | Adrenergic receptor agonist | ADRB2, ADRB1, ADRB3 |
| 515 | 92.56 | isoxsuprine | Adrenergic receptor agonist | ADRB2 |
| 553 | 92.25 | salmeterol | Adrenergic receptor agonist | ADRB2 |

ADRB2, beta2-adrenergic receptor; ADRB1, beta1-adrenergic receptor; ADRA1A, alpha1a-adrenergic receptor; ADRB3, beta3-adrenergic receptor.

Also, when CLUE was queried for connectivity with dopamine receptor agonists, 5 had high scores in relation to salbutamol; this included dopamine itself (Table 3). In view of the apparent associations between epinephrine and PD, it would appear to be important that there is an overlap of agonist activity of epinephrine on β2AR and dopamine receptors [43,44].

**Table 3.** Gene-expression connectivity scores of dopamine receptor agonists in relation to salbutamol.

| Rank | Score | Name | Description | Target |
| --- | --- | --- | --- | --- |
| 8 | 99.65 | SKF-77434 | Dopamine receptor agonist | DRD1 |
| 51 | 98.70 | fenoldopam | Dopamine receptor agonist | DRD1, ADRA1A, ADRA1B, ADRA1D, ADRA2A, ADRA2B, ADRA2C, DRD4, DRD5 |
| 52 | 98.66 | pramipexole | Dopamine receptor agonist | DRD3, DRD2, ADRA2A, ADRA2B, ADRA2C, DRD1, DRD4, DRD5, HTR1A, HTR1B, HTR1D, HTR2A, HTR2B, HTR2C |
| 335 | 93.45 | RO-10-5824 | Dopamine receptor agonist | DRD4 |
| 385 | 92.63 | dopamine | Dopamine receptor agonist | DRD2, DRD1, DRD3, DRD4, DRD5, DBH, TR1A, HTR7, SLC6A2, SLC6A3, SLC6A4 |

DRD1, dopamine receptor D1; DRD2, dopamine receptor D2; DRD3, dopamine receptor D3; DRD4, dopamine receptor D4; DRD5, dopamine receptor D5; ADRA1A, alpha1a-adrenergic receptor; ADRA1B, alpha1b-adrenergic receptor; ADRA1D, alpha1d-adrenergic receptor; ADRA2A, alpha2a-adrenergic receptor; ADRA2B, alpha2b-adrenergic receptor; ADRA2C, alpha2c-adrenergic receptor; HTR1A, HTR1B, HTR1D, HTR2A, HTR2B, HTR2C, serotonin receptors 1A, 1B, 1D, 2A, 2B, 2C; DBH, dopamine beta-hydroxylase; TR1A, T regulatory Type 1 cells; HTR7, serotonin 7 receptor; SLC6A2, norepinephrine transporter; SLC6A3, sodium dependent dopamine transporter; SLC6A4. serotonin transporter.

The potential anti-inflammatory importance of β2-adrenergic agonists was first observed in the current study with the observation of the marked similarity of the gene expression signatures of epinephrine to those of glucocorticoid receptor (GR) agonists. A portion of the heat map with the highest tau scores when epinephrine was modeled in CLUE already showed the predominance of GR agonists with high tau scores for the top matches in the database; the protein targets of the matching compounds were also replete with the codes, NRC31, glucocorticoid receptor, and NRC32, mineralocorticoid receptor (Fig. 1). In comparison, norepinephrine, which has approximately equal β1-adrenergic activity as epinephrine but has little or no β2 activity, showed essentially no connectivity to GR agonists in a comparable portion of its heatmap (Fig. 2). More than 60% of the gene expression signatures with connectivity to epinephrine in Fig. 1 were GR agonists while there was only 1 (0.025%) GR agonist that matched with a high tau score with norepinephrine (Fig. 2.).

**Figure 1.**
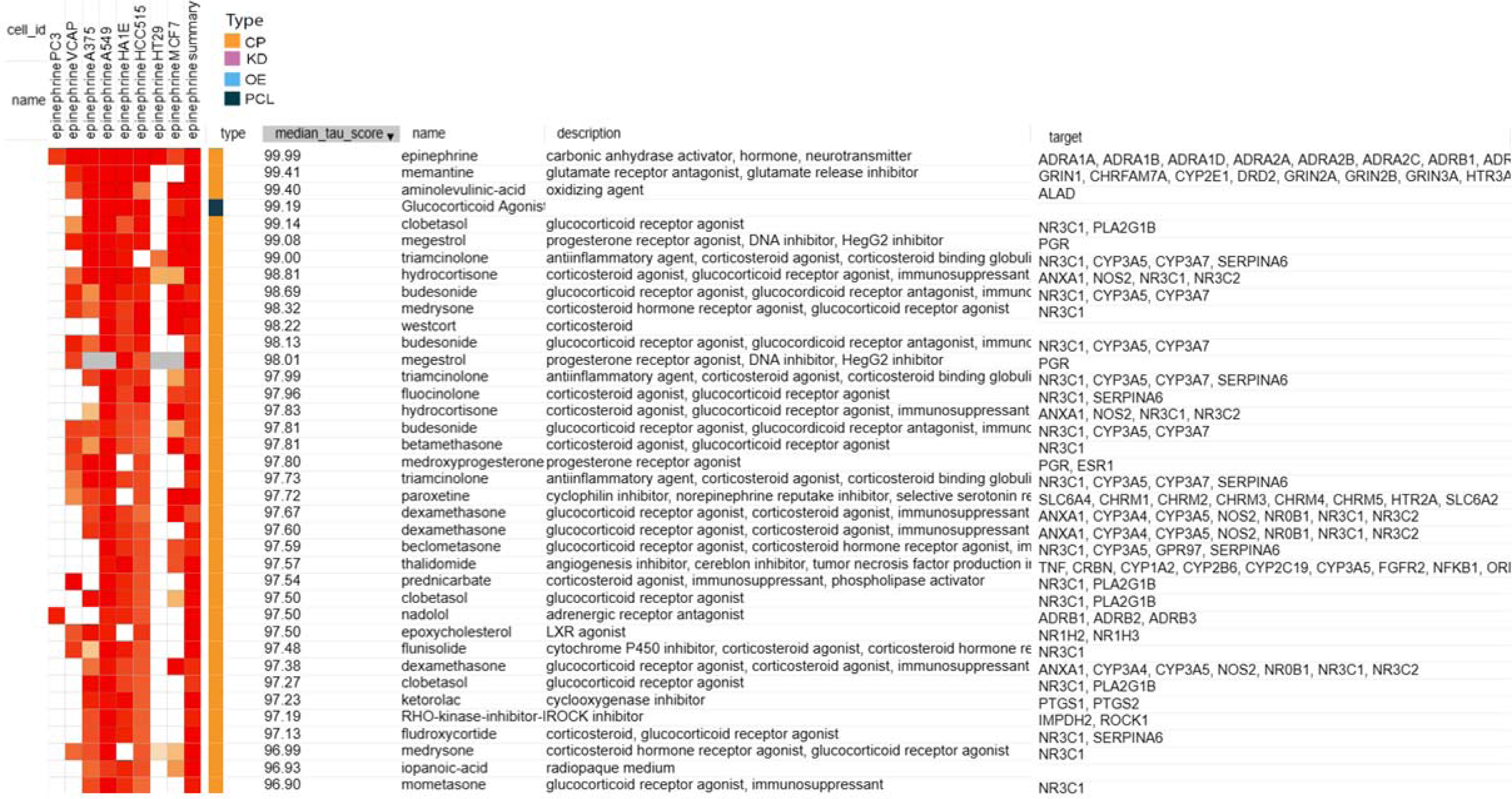
HeatMap of association streugths (median tau score) between gene expression signatures of epinephrine in 8 malignant cell cultures and those for all entries in the Touchstone database, with targets. Chemical compound (CP); Gene knock-down (KD); Gene over-expression (OE); and perturbagen class (PCL).

**Figure 2.**
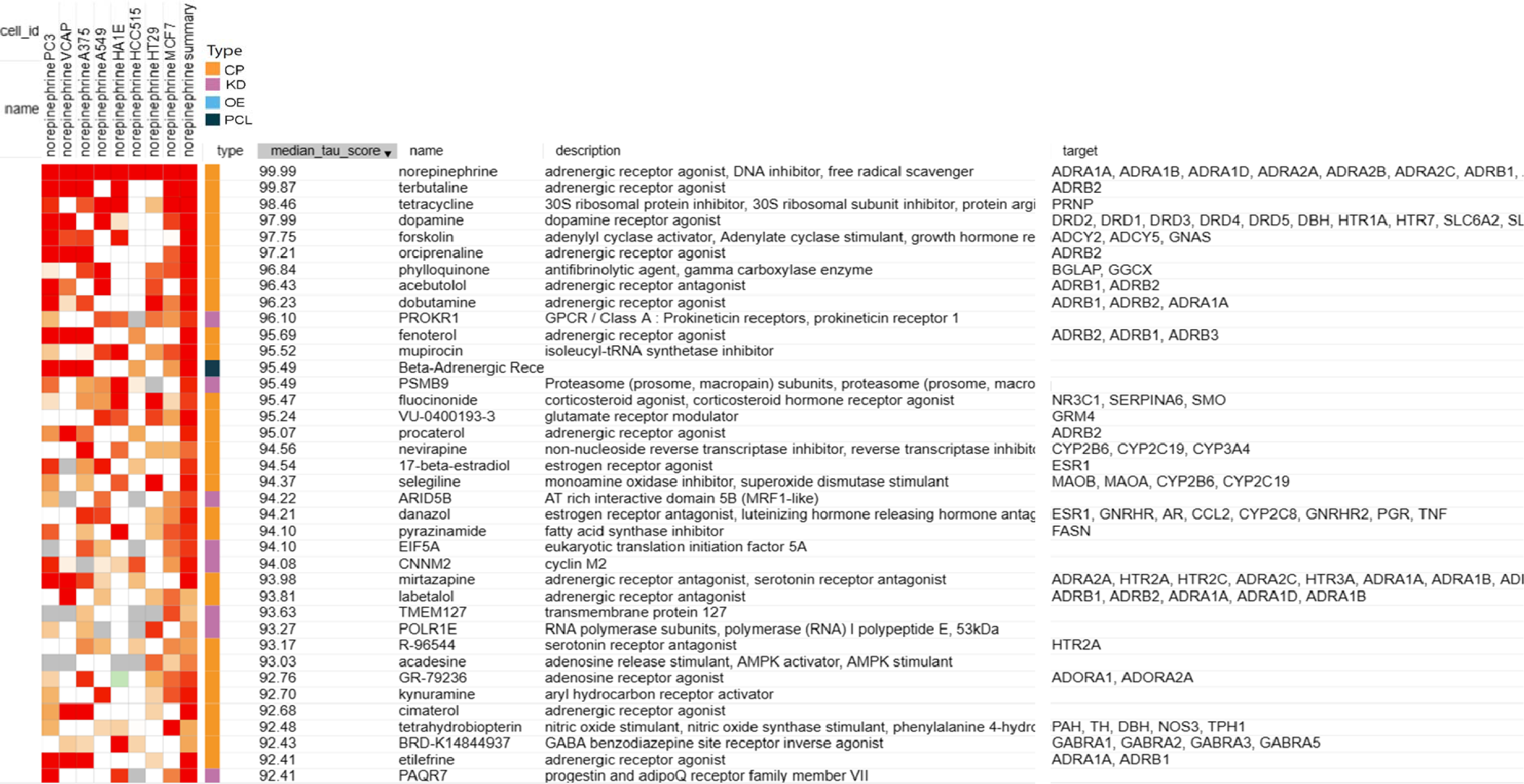
HeatMap of association strengths (median tau score) between gene expression signatures of noreplnephrlne In 8 malignant cell cultures and those for all entries In the Touchstone database. Chemlcal/pharmacologlc agent (CP); Gene knock-down (KD); Gene over-expression(OE); and pertu.rbagen class (PCL).

The detailed analysis outputs from the CLUE platform added further evidence of the strong association of epinephrine with drugs that have GR receptor agonist activity. A table is produced that shows the gene enrichment scores for CMap classes in the Touchstone database (Table 4). A high gene enrichment score is noted for the 44 GR agonists in this CMap class. As would be expected, there is also a high enrichment score for the 11 beta-adrenergic receptor agonists CMap class (Table 4). And, consistent with the almost complete lack of GR receptor agonists in the norepinephrine heatmap (Fig. 2), no connections with the GR CMap class are found in the comparable CMap class output for norepinephrine (Table 5). However, the general category of beta-adrenergic receptor agonists is represented (Table 5) because epinephrine and norepinephrine have equal β1-adrenergic potency.

**Table 4.** CMap Class Connectivity: EPINEPHRINE.

| **Perturbagen Class (PCL)** | **Group Enrichment Score** |
| --- | --- |
| Progesterone receptor agonist x 12 | 100.00 |
| Glucocorticoid receptor agonist x 44 | 99.67 |
| Aromatase inhibitor x 3 | 97.00 |
| Beta-adrenergic receptor agonist x 11 | 96.12 |
| FLT3 inhibitor x 6 | 95.46 |
| Progesterone receptor antagonist x 5 | 95.13 |
| Leucine rich repeat kinase inhibitor x 3 | 94.15 |
| Bromodomain Inhibitor x 6 | 93.59 |
| DNA synthesis inhibitor x 3 | 92.13 |
| VEGFR inhibitor x 13 | 91.84 |
| RAF inhibitor x 9 | 91.19 |

“Sets of compound perturbagens with enrichment scores above 90 (similar) and below -90 (opposing)” in relation to their correspondence with the gene expression signatures of epinephrine.

**Table 5.** CMap Class Connectivity: NOREPINEPHRINE.

| <u>Perturbagen Class (PCL)</u> | <u>Group Enrichment Score</u> |
| --- | --- |
| Beta-adrenergic receptor agonist x 11 | 99.95 |
| Adenosine receptor agonist x 6 | 99.58 |
| Reverse transcriptase inhibitor x 4 | 99.27 |
| Bacterial 30S ribosomal subunit inhibitor x 5 | 99.26 |
| GABA receptor antagonist x 5 | 95.90 |
| Sigma receptor antagonist x 3 | 95.58 |
| Bile acid x 4 | 93.38 |
| PPAR receptor agonist x 16 | 92.04 |
| Benzodiazepine receptor agonist x 4 | 90.55 |
| FXR antagonist x 3 | 90.27 |
| Estrogen receptor agonist x 17 | 90.13 |

“Sets of compound perturbagens with enrichment scores above 90 (similar) and below -90 (opposing)” in relation to their correspondence with the gene expression signatures of norepinephrine.

Still further evidence that an association exists between GR receptor agonist activity and the neurotransmitter, epinephrine, and that this is unique to this agent that has β2-adrenergic activity and not shared by norepinephrine, is seen in a comparison of the detailed analyses associated with the CMap class score differences. The strength of associations in gene signatures with GR receptor agonists for epinephrine and norepinephrine are presented in Tables 6 and 7, respectively. Of the 44 glucocorticoids in its CMap class, all have gene signature connectivity scores above 90.0 in relation to epinephrine (Table 6), while only 7 have scores above 90.0 in connectivity with norepinephrine and some have negative (opposing) connectivity (Table 7). These rankings are presented graphically in Fig. 3.

**Fig 3.**
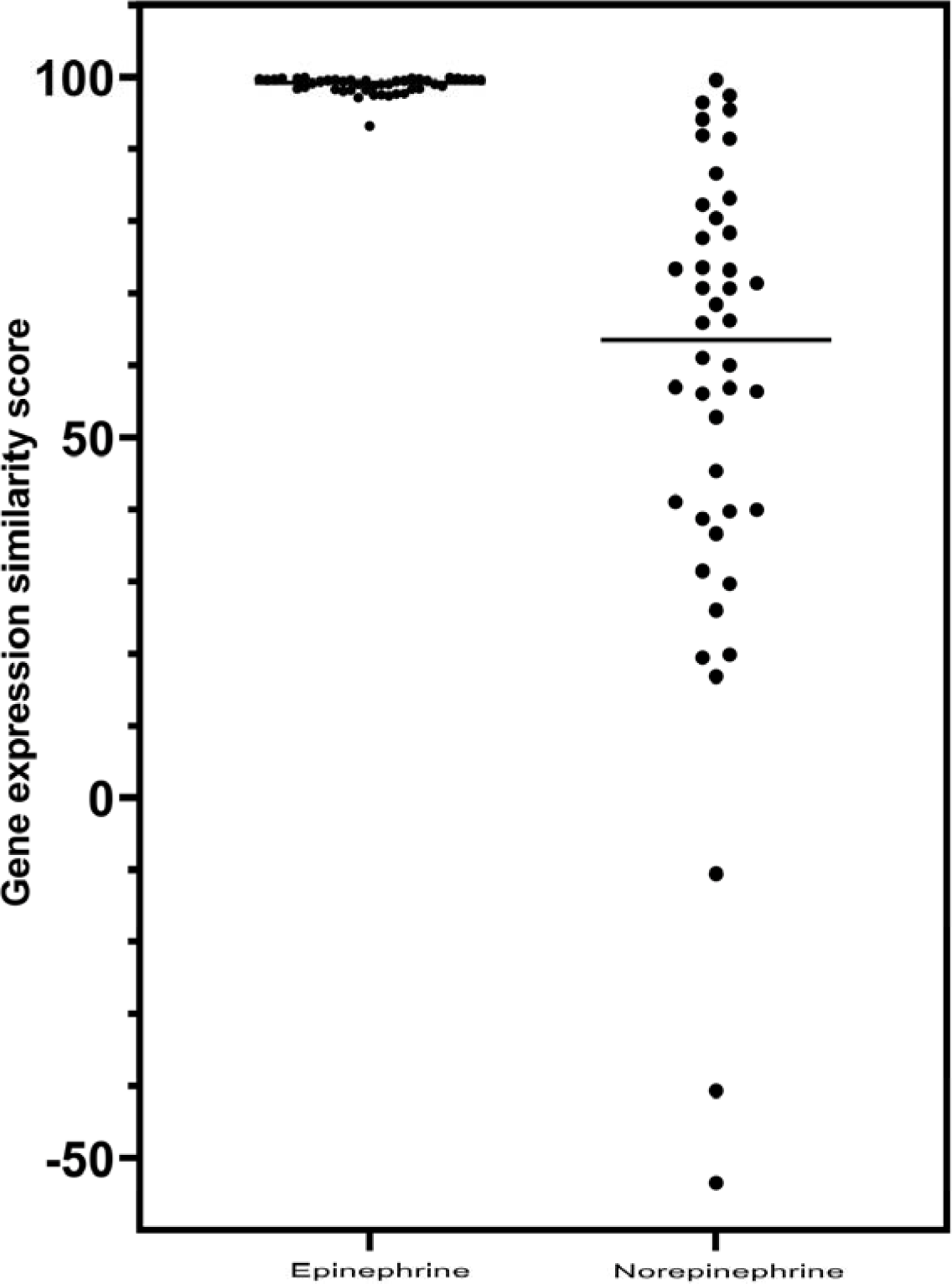
Comparison of epinephrine and norepinephrine gene-expression scores as glucocorticoid receptor agonists. The horizontal lines are the median values for the individual distributions. Mann-Whitney analysis of the distributions showed a statistically significant difference (p <0.0001), with strong association for epinephrine with gene-expression similarity to glucocorticoid agonists.

**Table 6.** Gene Expression-Connectivity Scores for EPINEPHRINE vs GR Agonists.

| <u>Rank</u> | <u>Score</u> | <u>Name</u> | <u>Description</u> |
| --- | --- | --- | --- |
| 4 | 99.89 | hydrocortisone | Glucocorticoid receptor agonist |
| 5 | 99.89 | betamethasone | Glucocorticoid receptor agonist |
| 7 | 99.79 | budesonide | Glucocorticoid receptor agonist |
| 8 | 99.79 | flunisolide | Cytochrome P450 inhibitor |
| 9 | 99.75 | mometasone | Glucocorticoid receptor agonist |
| 11 | 99.75 | fluocinonide | Glucocorticoid receptor agonist |
| 13 | 99.68 | triamcinolone | Glucocorticoid receptor agonist |
| 14 | 99.68 | fludroxycortide | Glucocorticoid receptor agonist |
| 19 | 99.62 | clobetasol | Glucocorticoid receptor agonist |
| 21 | 99.61 | triamcinolone | Glucocorticoid receptor agonist |
| 22 | 99.61 | flumetasone | Glucocorticoid receptor agonist |
| 23 | 99.58 | fluticasone | Glucocorticoid receptor agonist |
| 24 | 99.58 | fluorometholone | Glucocorticoid receptor agonist |
| 26 | 99.54 | dexamethasone | Glucocorticoid receptor agonist |
| 27 | 99.52 | prednicarbate | Phospholipase activator |
| 28 | 99.51 | isoflupredone | Glucocorticoid receptor agonist |
| 29 | 99.47 | hydrocortisone | Glucocorticoid receptor agonist |
| 30 | 99.44 | diflorasone | Corticosteroid agonist |
| 31 | 99.44 | dexamethasone | Glucocorticoid receptor agonist |
| 32 | 99.4 | medrysone | Glucocorticoid receptor agonist |
| 33 | 99.37 | prednisolone | Glucocorticoid receptor agonist |
| 34 | 99.33 | halcinonide | Glucocorticoid receptor agonist |
| 37 | 99.12 | betamethasone | Glucocorticoid receptor agonist |
| 39 | 99.05 | beclometasone | Glucocorticoid receptor agonist |
| 40 | 99.05 | fluocinolone | Glucocorticoid receptor agonist |
| 42 | 99.05 | hydrocortisone | Glucocorticoid receptor agonist |
| 43 | 99.01 | hydrocortisone | Glucocorticoid receptor agonist |
| 45 | 98.77 | clocortolone | Glucocorticoid receptor agonist |
| 46 | 98.77 | beclometasone | Glucocorticoid receptor agonist |
| 48 | 98.6 | westcort | Glucocorticoid receptor agonist |
| 51 | 98.4 | prednisolone | Glucocorticoid receptor agonist |
| 52 | 98.38 | fluticasone | Glucocorticoid receptor agonist |
| 55 | 98.34 | hydrocortisone | Glucocorticoid receptor agonist |
| 56 | 98.32 | rimexolone | Glucocorticoid receptor agonist |
| 58 | 98.17 | fludrocortisone | Glucocorticoid receptor agonist |
| 59 | 98.17 | amcinonide | Glucocorticoid receptor agonist |
| 61 | 98.03 | methylprednisolone | Glucocorticoid receptor agonist |
| 67 | 97.71 | desoximetasone | Glucocorticoid receptor agonist |
| 75 | 97.67 | halometasone | Glucocorticoid receptor agonist |
| 88 | 97.53 | loteprednol | Glucocorticoid receptor agonist |
| 96 | 97.5 | depomedrol | Glucocorticoid receptor agonist |
| 117 | 97.39 | prednisolone | Glucocorticoid receptor agonist |
| 137 | 97.15 | fluocinonide | Glucocorticoid receptor agonist |
| 632 | 93.24 | alclometasone | Glucocorticoid receptor agonist |

**Table 7.** Gene-Expression Connectivity Scores for NOREPINEPHRINE vs GR agonists.

| <u>Rank</u> | <u>Score</u> | <u>Name</u> | <u>Description</u> |
| --- | --- | --- | --- |
| 23 | 99.55 | fluocinonide | Glucocorticoid receptor agonist |
| 244 | 97.47 | depomedrol | Glucocorticoid receptor agonist |
| 376 | 96.44 | beclometasone | Glucocorticoid receptor agonist |
| 471 | 95.46 | fluocinolone | Glucocorticoid receptor agonist |
| 615 | 94.13 | prednisolone | Glucocorticoid receptor agonist |
| 890 | 91.89 | triamcinolone | Glucocorticoid receptor agonist |
| 943 | 91.4 | amcinonide | Glucocorticoid receptor agonist |
| 1479 | 86.61 | rimexolone | Glucocorticoid receptor agonist |
| 1810 | 83.17 | hydrocortisone | Glucocorticoid receptor agonist |
| 1959 | 82.28 | halometasone | Glucocorticoid receptor agonist |
| 2159 | 80.37 | beclometasone | Glucocorticoid receptor agonist |
| 2349 | 78.38 | triamcinolone | Glucocorticoid receptor agonist |
| 2445 | 77.7 | prednisolone | Glucocorticoid receptor agonist |
| 2855 | 73.57 | betamethasone | Glucocorticoid receptor agonist |
| 2864 | 73.36 | hydrocortisone | Glucocorticoid receptor agonist |
| 2877 | 73.18 | alclometasone | Glucocorticoid receptor agonist |
| 3084 | 71.41 | westcort | Glucocorticoid receptor agonist |
| 3155 | 70.7 | fluticasone | Glucocorticoid receptor agonist |
| 3160 | 70.64 | loteprednol | Glucocorticoid receptor agonist |
| 3348 | 68.45 | dexamethasone | Glucocorticoid receptor agonist |
| 3561 | 66.22 | fluocinonide | Glucocorticoid receptor agonist |
| 3591 | 65.9 | desoximetasone | Glucocorticoid receptor agonist |
| 4051 | 61.03 | betamethasone | Glucocorticoid receptor agonist |
| 4147 | 60 | prednicarbate | Phospholipase activator |
| 4393 | 56.96 | clocortolone | Glucocorticoid receptor agonist |
| 4404 | 56.86 | hydrocortisone | Glucocorticoid receptor agonist |
| 4465 | 56.37 | fluorometholone | Glucocorticoid receptor agonist |
| 4489 | 56.05 | flunisolide | Cytochrome P450 inhibitor |
| 4701 | 52.82 | mometasone | Glucocorticoid receptor agonist |
| 5218 | 45.35 | dexamethasone | Glucocorticoid receptor agonist |
| 5512 | 41.08 | prednisolone | Glucocorticoid receptor agonist |
| 5594 | 39.97 | fludroxycortide | Glucocorticoid receptor agonist |
| 5618 | 39.72 | methylprednisolone | Glucocorticoid receptor agonist |
| 5696 | 38.68 | fluticasone | Glucocorticoid receptor agonist |
| 5837 | 36.6 | hydrocortisone | Glucocorticoid receptor agonist |
| 6223 | 31.46 | medrysone | Glucocorticoid receptor agonist |
| 6329 | 29.69 | budesonide | Glucocorticoid receptor agonist |
| 6560 | 25.98 | isoflupredone | Glucocorticoid receptor agonist |
| 6962 | 19.87 | clobetasol | Glucocorticoid receptor agonist |
| 7001 | 19.45 | fludrocortisone | Glucocorticoid receptor agonist |
| 7129 | 16.81 | halcinonide | Glucocorticoid receptor agonist |
| 8126 | -10.56 | flumetasone | Glucocorticoid receptor agonist |
| 8351 | -40.7 | hydrocortisone | Glucocorticoid receptor agonist |
| 8433 | -53.42 | diflorasone | Corticosteroid agonist |

It is of particular interest, and unexpected, that the perturbagen class output for epinephrine (Table 4) included leucine rich repeat kinase (LRRK) inhibitors as showing gene-expression connectivity to epinephrine with a strong positive score. Among the agents in the CLUE database, LRRK2 inhibitors, XMD-1150, Broad Institute ID (BRD)-K01436366 and XMD-885 (BRD-K64857848), ranked with connectivity scores to epinephrine of 90.78 and 90.47, respectively. LRRK2 inhibitors are currently a major focus as potential treatments for PD [45–48]. Table 4 also includes a positive score for gene-expression connectivity with epinephrine for vascular endothelial growth factor receptor (VEGFR) inhibitors; this will be commented upon below. Quite remarkably, the CLUE output in Table 4 also includes a positive score for connectivity of epinephrine with rapidly accelerated fibrosarcoma (RAF) inhibitors. The proto-oncogene, *B-RAF*, is part of the pathway of *RAF* activation of many cancers [49]. Although this could be of considerable interest in relation to possible anti-neoplastic activity of β2AR agonists, it was not pursued in the current study.

It can be noted that the positive rankings for LRRK2, VEGFR, and RAF inhibitors with epinephrine were not identified with gene expression connectivity to norepinephrine (Table 5), which is consistent with the association of these properties with β2-adrenergic activity.

The CLUE analyses of a marked difference between the associations of epinephrine and norepinephrine with gene signatures that are predictive of GR agonist activity are parallel with the established biological differences between the agonist activities of epinephrine and norepinephrine (Table 8). Epinephrine and norepinephrine have essentially equal α1-, α2- and β1-adrenergic agonist activity, but epinephrine is substantially more active than norepinephrine as a β2-adrenergic agonist. The evidence outlined above that supports an anti-inflammatory activity of β2-adrenergic agonists, and a potentially therapeutic effect in relation to the neuroinflammation aspect of PD, is a major focus of these investigations. Also, the fact that GR agonists are widely recognized for their anti-inflammatory activity adds to the plausibility of these associations with β2-adrenergic activity.

**Table 8.** Modified from: Goodman and Gilman’s The Pharmacological Basis of Therapeutics, 9^th^ Edition, 1996. Joel G. Hardman and Lee E. Limbird, Eds. McGraw-Hill, New York. Chapter 6, Neurotransmission, Page 125 (Table 6-3).

**Characteristics of Subtypes of Adrenergic Receptors<sup>1</sup>**
| Receptor | Agonists | Antagonists | Tissue | Responses |
| --- | --- | --- | --- | --- |
| Alpha1 | Epi ≥NE>>Iso<br>Drug example, phenylephrine | Prazosin | Vascular smooth muscle | Contraction |
|  |  |  | Genitourinary smooth muscle | Contraction |
|  |  |  | Liver | Glycogenolysis; gluconeogenesis |
|  |  |  | Intestinal smooth muscle | Hyperpolarization and relaxation |
|  |  |  | Heart | Increased contractile force, arrhythmias |
| Alpha2 | Epi ≥NE>>Iso<br>Drug example, Clonidine | Yohimbine | Pancreatic islet (β cells) | Decreased insulin secretion |
|  |  |  | Platelets | Aggregation |
|  |  |  | Nerve terminals | Decreased release of NE |
|  |  |  | Vascular Smooth Muscle | Contraction |
| Beta1 | Iso>>Epi=NE<br>Drug example, Dobutamine | Metoprolol | Heart | Increased force and rate of contraction and AV nodal conduction velocity |
|  |  |  | Juxtaglomerular cell | Increased renin secretion |
| Beta2 | Iso>>Epi >>>NE<br>Drug example, Terbutaline | ICI 118551 (Inverse agonist) | Smooth muscle (vascular, bronchial, gastrointestinal, and genitourinary) | Relaxation |
|  |  |  | Skeletal muscle | Glycogenolysis; uptake of K<sup>+</sup> |
|  |  |  | Liver | Glycogenolysis; gluconeogenesis |
| Beta3 | Iso=NE>Epi | ICI 118551 | Adipose tissue | Lipolysis |

<sup>1</sup>This table provides examples of drugs that act on adrenergic receptors and of the location of subtypes of adrenergic receptors. Abbreviations: epinephrine (Epi); norepinephrine (NE); isoproterenol (Iso). Epi ≥NE>>Iso (Epi equal to/slightly greater than NE, which is greater than Iso); Iso>>Epi=NE (Iso greater than Epi, which is equal to NE); Iso>>Epi>>>NE (Iso greater than Epi, which is much greater than NE); Iso=NE>Epi (Iso equal to NE, which is slightly greater than Epi).

When the natural endogenous human glucocorticoid, cortisol (hydrocortisone), was profiled in CLUE, as expected, it showed strong gene signature connectivity with the synthetic glucocorticoid drugs in the CLUE CMap class (Table 9). Interestingly, it also ranked the beta-adrenergic receptor agonist class with a high enrichment score, similar to that for epinephrine (Table 4). The detailed analyses also demonstrated that the 44 synthetic glucocorticoids in the CMap class all had gene signature connectivity scores above 90.0 with hydrocortisone (Table 10); again, this is similar to the finding with epinephrine (Table 6).

**Table 9.** CMap Class Connectivity: HYDROCORTISONE.

| **Perturbagen Class (PCL)** | **Group Enrichment Score** |
| --- | --- |
| Glucocorticoid receptor agonist x 44 | 99.89 |
| Cyclooxygenase inhibitor x 6 | 97.34 |
| Beta-adrenergic receptor agonist x 11 | 95.88 |
| Adenosine receptor agonist x 6 | 95.49 |
| Na-K-Cl transporter inhibitor x 3 | 94.89 |
| Rho associated kinase inhibitor x 4 | 93.79 |
| MEK inhibitor x 8 | 92.61 |

“Sets of compound perturbagens with enrichment scores above 90 (similar) and below -90 (opposing)” in relation to their correspondence with the gene expression signatures of hydrocortisone.

**Table 10.** Gene-expression connectivity scores for HYDROCORTISONE vs GR agonists.

| **Rank** | **Score** | **Name** | **Description** |
| --- | --- | --- | --- |
| 1 | 99.99 | hydrocortisone | Glucocorticoid receptor agonist |
| 2 | 99.93 | triamcinolone | Glucocorticoid receptor agonist |
| 3 | 99.93 | budesonide | Glucocorticoid receptor agonist |
| 4 | 99.93 | fluocinonide | Glucocorticoid receptor agonist |
| 5 | 99.93 | fluorometholone | Glucocorticoid receptor agonist |
| 7 | 99.72 | mometasone | Glucocorticoid receptor agonist |
| 8 | 99.72 | fluticasone | Glucocorticoid receptor agonist |
| 9 | 99.72 | fludrocortisone | Glucocorticoid receptor agonist |
| 11 | 99.61 | betamethasone | Glucocorticoid receptor agonist |
| 12 | 99.61 | betamethasone | Glucocorticoid receptor agonist |
| 13 | 99.58 | clocortolone | Glucocorticoid receptor agonist |
| 14 | 99.58 | flunisolide | Cytochrome P450 inhibitor |
| 15 | 99.58 | dexamethasone | Glucocorticoid receptor agonist |
| 16 | 99.58 | clobetasol | Glucocorticoid receptor agonist |
| 17 | 99.54 | isoflupredone | Glucocorticoid receptor agonist |
| 20 | 99.37 | fludroxycortide | Glucocorticoid receptor agonist |
| 21 | 99.33 | prednisolone | Glucocorticoid receptor agonist |
| 22 | 99.3 | beclomethasone | Glucocorticoid receptor agonist |
| 23 | 99.3 | fluticasone | Glucocorticoid receptor agonist |
| 24 | 99.3 | alclometasone | Glucocorticoid receptor agonist |
| 26 | 99.3 | hydrocortisone | Glucocorticoid receptor agonist |
| 27 | 99.26 | fluocinolone | Glucocorticoid receptor agonist |
| 28 | 99.25 | flumetasone | Glucocorticoid receptor agonist |
| 30 | 99.16 | rimexolone | Glucocorticoid receptor agonist |
| 32 | 99.06 | loteprednol | Glucocorticoid receptor agonist |
| 34 | 98.91 | hydrocortisone | Glucocorticoid receptor agonist |
| 35 | 98.8 | halcinonide | Glucocorticoid receptor agonist |
| 36 | 98.66 | triamcinolone | Glucocorticoid receptor agonist |
| 37 | 98.63 | dexamethasone | Glucocorticoid receptor agonist |
| 38 | 98.59 | beclomethasone | Glucocorticoid receptor agonist |
| 39 | 98.52 | prednicarbate | Phospholipase activator |
| 44 | 98.48 | prednisolone | Glucocorticoid receptor agonist |
| 45 | 98.48 | amcinonide | Glucocorticoid receptor agonist |
| 46 | 98.1 | halometasone | Glucocorticoid receptor agonist |
| 48 | 97.82 | methylprednisolone | Glucocorticoid receptor agonist |
| 50 | 97.68 | diflorasone | Corticosteroid agonist |
| 52 | 97.5 | hydrocortisone | Glucocorticoid receptor agonist |
| 53 | 97.5 | fluocinonide | Glucocorticoid receptor agonist |
| 55 | 97.36 | desoximetasone | Glucocorticoid receptor agonist |
| 58 | 97.33 | prednisolone | Glucocorticoid receptor agonist |
| 66 | 97.15 | depomedrol | Glucocorticoid receptor agonist |
| 83 | 96.69 | medrysone | Glucocorticoid receptor agonist |
| 141 | 95.45 | westcort | Glucocorticoid receptor agonist |
| 187 | 95.1 | hydrocortisone | Glucocorticoid receptor agonist |

To summarize, Fig. 4 presents the comparison between the gene expression signatures when CLUE is probed with epinephrine and hydrocortisone as INDEX compounds (i.e., Table 6 vs. Table 10 results).

**Fig 4.**
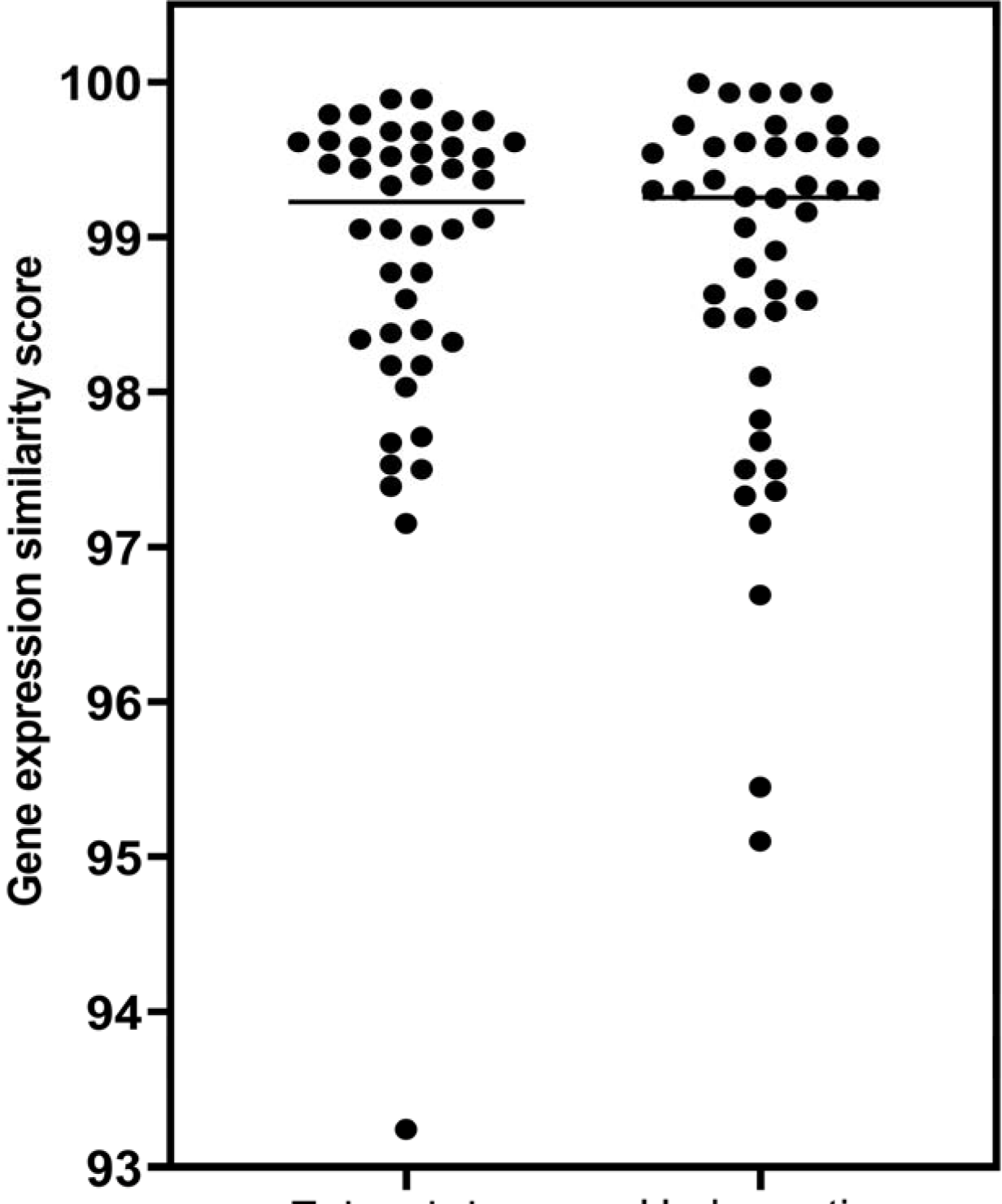
Comparison of epinephrine and hydrocortisone gene-expression scores as glucocorticoid receptor agonists. The horizontal lines are the median values for the individual distributions, which are essentially identical. Mann-Whitney analysis of the distributions showed no difference in gene-expression characteristics for epinephrine and hydrocortisone as glucocorticoid receptor agonists (p = 0.805).

The gene-product targets for the 44 glucocorticoid analyses in this CMap class are presented in S1 Table. All members of this class have the glucocorticoid receptor, NR3C1, as a target; some also have the mineralocorticoid receptor, NR3C2, as a target. Only the natural human glucocorticoid cortisol (hydrocortisone) and the synthetic drugs dexamethasone and amcinonide have the gene product ANXA1 as a target. Annexin A1, also termed lipocortin I, is the product of the gene *ANXA1,* and appears to primarily serve the anti-inflammatory effect of several glucocorticoids. [50] In addition to the gene-product targets for the glucocorticoids, S1 Table includes the Broad Institute ID numbers for each analysis of gene-expression in the database.

It should be noted that the CMap class output for hydrocortisone (Table 9) included a strong group score (92.61) for gene-expression connectivity to mitogen-activated protein kinase kinase (MEK) inhibitors. This kinase is on the pathway to signaling of malignant proliferation and survival, i.e., RAF > MEK > extracellular signal-related kinases (ERK). This is similar to the observations noted above for epinephrine (Table 4) where gene-expression connectivity was observed for inhibitory mediators of the anti-proliferative/anti-neoplastic effects of LRRK2, VEGFR, and RAF. In the case of the MEK inhibitors, the drug with the highest score for the class when probed separately in CLUE was selumetinib (96.51), which has proven to have a broad role as an anti-neoplastic agent [51, 52].

The striking similarities in the gene-expression signature connectivity for epinephrine and hydrocortisone drew attention to their known interrelationships in regard to synthesis of cAMP [53–55]. Perez [53] noted that α2 agonist activity mediated a decreased synthesis of cAMP. When the classical non-competitive, irreversible α2 antagonist, phenoxybenzamine, was modeled in CLUE, it also showed considerable gene-expression signature connectivity with GR receptor agonists. The gene enrichment score for phenoxybenzamine with the GR agonist CMap class is shown in Table 11.

**Table 11.** CMap Class Connectivity: PHENOXYBENZAMINE.

| **Perturbagen Class (PCL)** | **Group Enrichment Score** |
| --- | --- |
| Progesterone receptor agonist x 12 | 99.50 |
| Glucocorticoid receptor agonist x 44 | 98.78 |
| Aromatase inhibitor x 3 | 97.98 |
| Glycogen synthase kinase inhibitor x 11 | 97.63 |
| HSP inhibitor x 7 | 96.76 |
| EGFR inhibitor x 14 | 96.33 |
| PPAR receptor agonist x 16 | 95.40 |
| DNA synthesis inhibitor x 3 | 95.02 |
| PKC inhibitor x 7 | 93.75 |
| JAK inhibitor x 5 | 92.59 |
| HIV protease inhibitor x 4 | 92.34 |
| Cannabinoid receptor agonist x 8 | 91.13 |

“Sets of compound perturbagens with enrichment scores above 90 (similar) and below -90 (opposing)” in relation to their correspondence with the gene expression signatures of phenoxybenzamine.

**Table 12.**
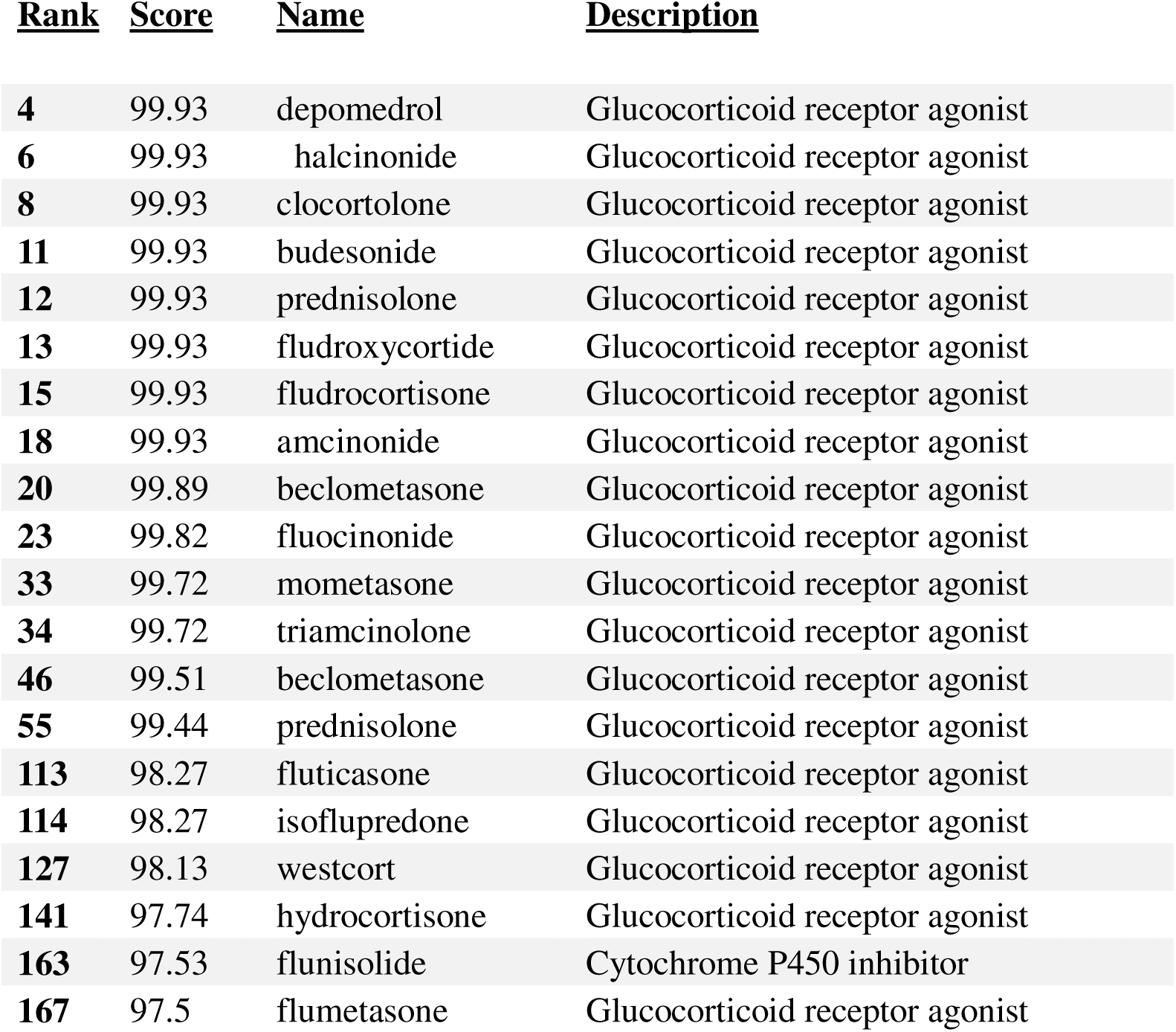
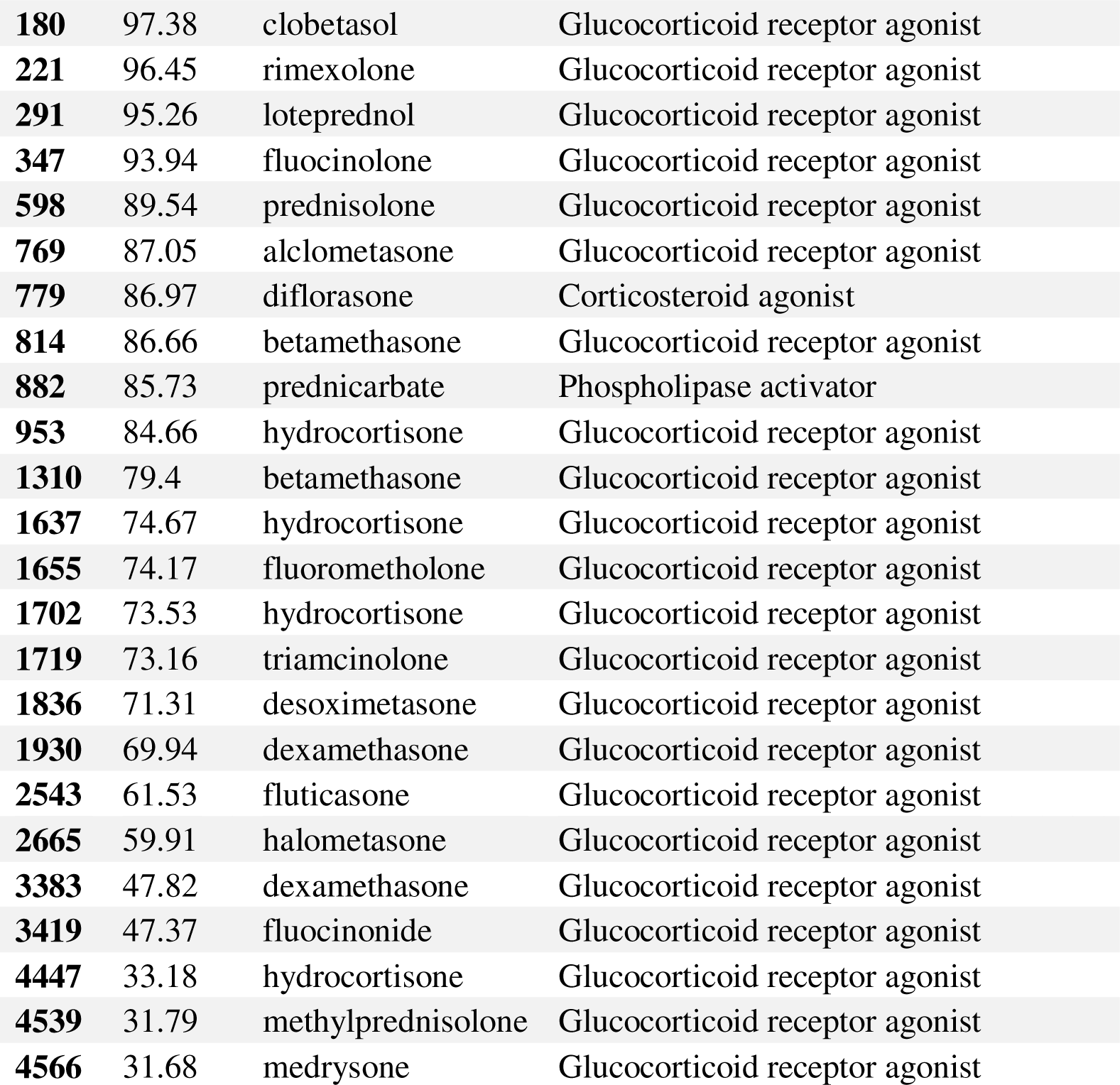
Gene-Expression Connectivity Scores for PHENOXYBENZAMINE vs GR agonists.

The strong gene-expression connectivity of phenoxybenzamine with GR agonist activity and the potential anti-neoplastic properties of phenoxybenzamine (Table 11) have been presented in previous reports [56, 57].

The associations between agents that increase cAMP levels and GR receptor agonist activity were investigated further by modeling two methylxanthine drugs, theophylline and caffeine, in CLUE. Theophylline has a long therapeutic history as a bronchodilator in the treatment of respiratory diseases. As discussed above, it does not directly increase synthesis of cAMP, but results in higher levels of cellular cAMP by inhibition of phosphodiesterase enzymes that metabolize cAMP. Caffeine has similar activity but is a much more potent CNS stimulant than a peripheral agonist.

The CMap class output scores from CLUE for theophylline are presented in Table 13. Gene-expression connectivity scores vs GR agonists are presented in S2 Table.

**Table 13.**
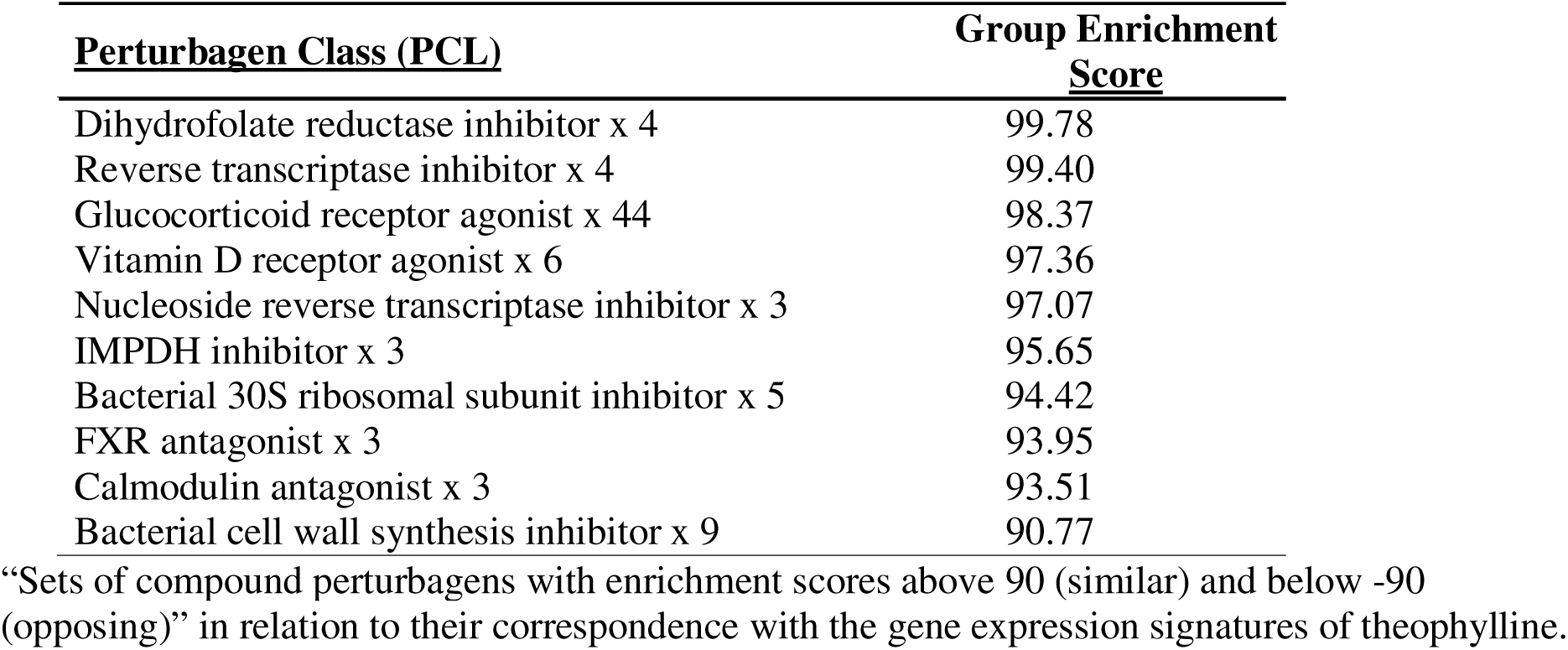
CMap Class Connectivity: THEOPHYLLINE.

The CMap class output scores from CLUE for caffeine are presented in Table 14. The GR agonist class scored substantially lower for caffeine than for theophylline. Gene-expression connectivity scores vs GR agonist are presented in S3 Table.

**Table 14.** CMap Class Connectivity: CAFFEINE.

| <u>Perturbagen Class (PCL)</u> | <u>Group Enrichment Score</u> |
| --- | --- |
| Bacterial DNA gyrase inhibitor x 5 | 95.54 |
| Aromatase inhibitor x 3 | 93.32 |
| Dihydrofolate reductase inhibitor x 4 | 91.76 |
| Cyclooxygenase inhibitor x 6 | 90.65 |
| Glucocorticoid receptor agonist x 44 | 90.24 |

“Sets of compound perturbagens with enrichment scores above 90 (similar) and below -90 (opposing)” in relation to their correspondence with the gene expression signatures of caffeine.

A summary of agents that have been investigated in relation to their gene-expression signatures with connectivity to GR agonists is presented in Figure 5.

**Fig 5.**
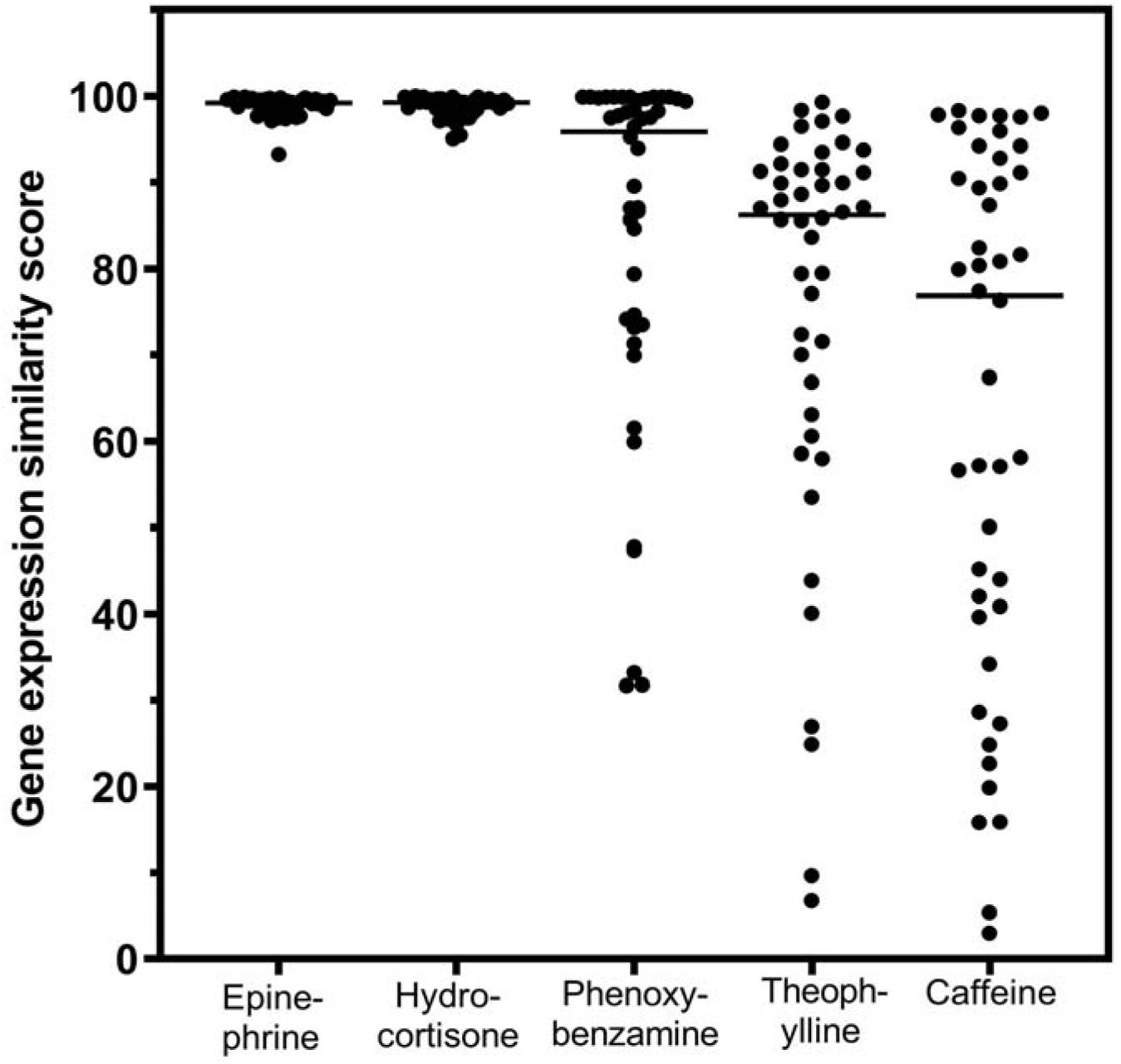
Comparison of beta2-adrenergic related agents and hydrocortisone for gene-expression connectivity scores as glucocorticoid receptor agonists. The horizontal lines are the median values for the individual distributions. The relative differences among the gene-expression scores can be appreciated visually. Caffeine was not studied further.

Epinephrine, a potent β2AR agonist has been presented as a prototype for drugs of this class in these studies because levalbuterol and arformoterol have not been profiled for gene-expression connectivity in CLUE. In July 2020, as our interest in levalbuterol advanced, we formally nominated it to the Broad Institute for gene-expression connectivity study in the CLUE database based on its apparent anti-inflammatory potency [7]; it remained on the queue for possible testing for several months but was never screened.

It would appear to be significant that the drugs noted in Figure 5 that have gene-expression signatures with connectivity to GR agonists are all capable of mediating an increase in cellular cAMP. Since increases in cAMP are a characteristic of drugs that mediate anti-inflammatory effects, it appears that this may represent a unifying hypothesis for this study.

### 3.2. CRS assays of agents with β2-adrenergic and related activities

The CRS assay was used to test the efficacy of the several β2-adrenergic related drugs discussed above for possible relationships between predictions generated in the CLUE genomic analyses and a biological test of anti-inflammatory activity in the humanized mouse assay. All animal assays were carried out in the Jackson Laboratories facility in Sacramento, California.

Several of the drugs of interest were tested for their ability to modulate cytokine release *in vivo* after the mice were challenged with injections of the monoclonal antibodies OKT3 or aCD28. The application of an *in vivo* model to simulate a clinical translation was considered a feature of these investigations. The drugs were administered in the drinking water. They are all water soluble and are known to have good bioavailability by the oral route. Doses were calculated from the standard human clinical range of mg/kg body weight per day. The FDA factor of 12.3 was applied for conversion from human to mouse mg/kg dose. A mouse body weight of 20 grams and an average intake of drinking water of 4 ml/day were assumed. The resulting doses were as follows: Albuterol sulfate, 8 µg/ml drinking water; levalbuterol HCl, 4 µg/ml; phenoxybenzamine HCl, 5 µg/ml; theophylline, 400 µg/ml; and arformoterol tartrate, 0.008 µg/ml.

Mice treated with levalbuterol showed a statistically significant lower body weight loss than the control group (1% sucrose) when CRS was induced with OKT3 (Figure 6).

**Fig 6.**
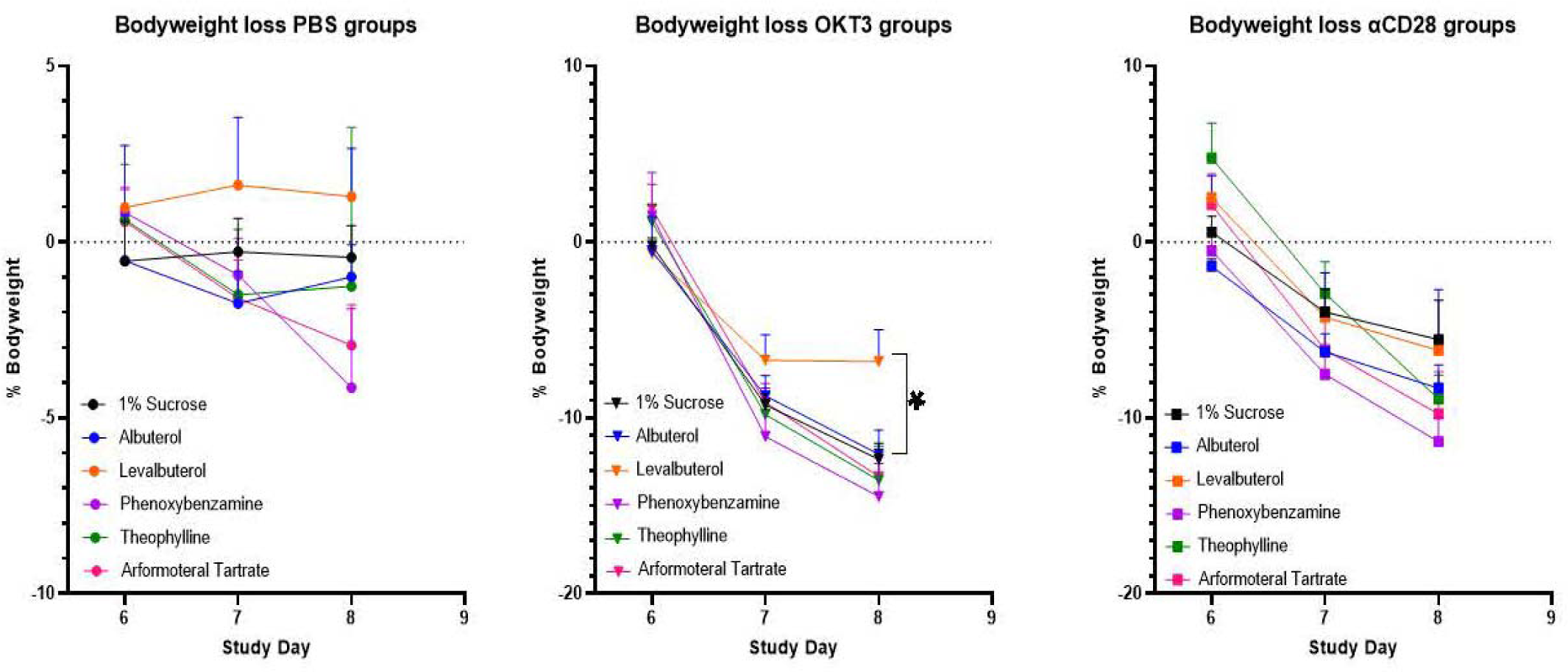
Percent body weight loss calculated from each animal’s initial body weight at Study Day 0. Body weight loss was statistically significantly less for the group that was pretreated with levalbuterol and challenged with the monoclonal antibody, OKT3.

There were no statistically significant differences in the clinical scoring between any of the groups and the control group (Figure 7). All 90 animals included in the study survived to the time of blood collection by cardiac puncture on Study Day 9, 72 hours after toxic induction with OKT3 or aCD28. This would suggest that the doses translated from human to mouse were reasonably appropriate.

**Fig 7.**
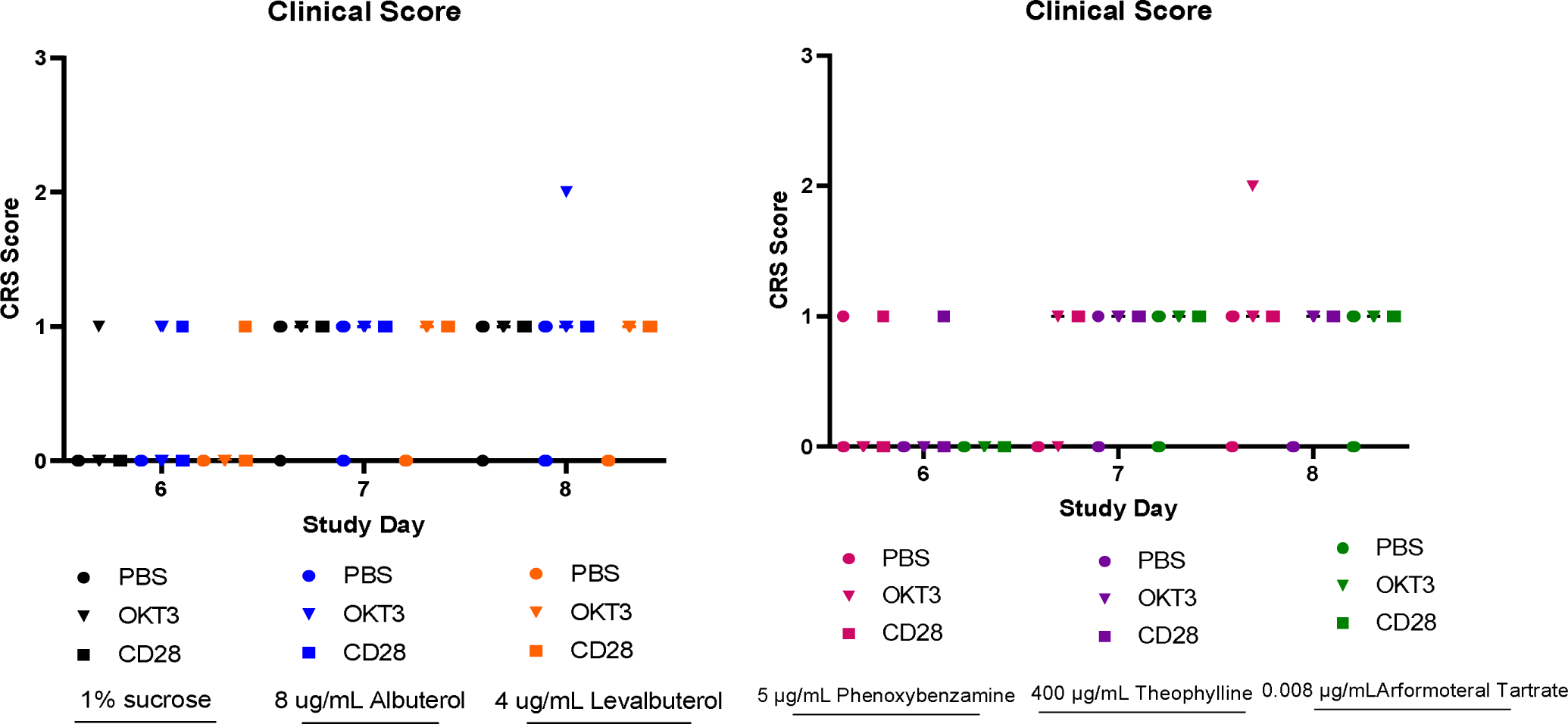
CRS clinical scores for the period following toxic induction of cytokine release with OKT3 or aCD28. There were no statistically significant differences in the clinical scores that were attributable to the pretreatments; the doses of pretreatments were calculated from typical human doses that were translated using FDA factors.

Water consumption in both the OKT3 and anti-CD28 groups peaked on Study Day 6 before toxicity was induced and declined progressively to Study Day 9 (Figure 8). Analysis of the data was limited since water consumption was made by the cage for each group of five mice. However, one individual comparison (Wilcoxon signed rank test) for water consumption for levalbuterol vs. sucrose control in the OKT3 groups did trend for a statistical advantage favoring more water intake for levalbuterol (p = 0.068). (Data for theophylline groups were accidentally not recorded.)

**Fig 8.**
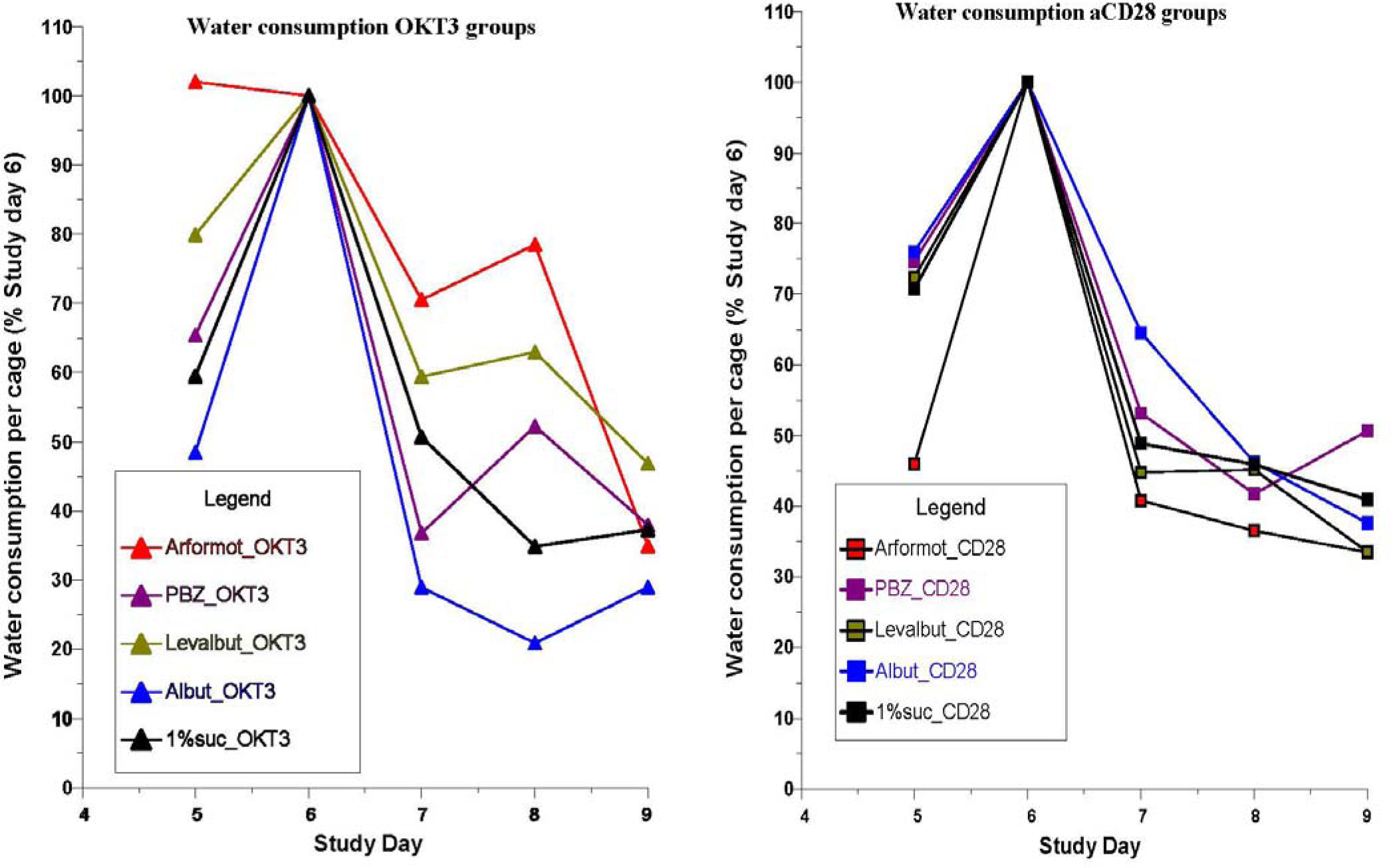
Water consumption recorded per cage of 5 mice as a percentage of the intake on Study Day 6 prior to the induction challenges with OKT3 and anti-CD28. The toxic antibody challenges to induce cytokine release caused marked decreases in water consumption in all of the groups. Since the pretreatment drugs were administered in the drinking water, it would be expected that drug delivery was compromised.

Analyses of the effects of the beta2 adrenergic agonists and related compounds on the release of cytokines and chemokines 6 hours after induction with OKT3 in the CRS assay demonstrated the following primary outcomes: Figure 9 presents the results for the inhibition of the release of the eosinophil-selective chemokine, eotaxin, which is responsible for a number of eosinophilic disorders, and the angiogenic proto-oncogene, vascular endothelial growth factor (VEGF), by several of the β2-adrenergic related drugs. Albuterol, the racemic mixture of the levo and dextro isomers, was not different from the sucrose control in any of the assays. It should be noted that albuterol was administered at twice the concentration of levalbuterol, therefore the mice received the same dose of the levo isomer as those receiving the pure levalbuterol. However, as noted above, the dextro isomer is pro-inflammatory and interferes with the anti-inflammatory activity of the levo isomer. Levalbuterol also inhibited the release of the pro-inflammatory cytokine, IL-13 with OKT3 induction (Figure 9). The relationships between the treatment drugs and their gene-expression connectivity to glucocorticoid receptor agonist activity is also included in Figure 9.

**Fig 9.**
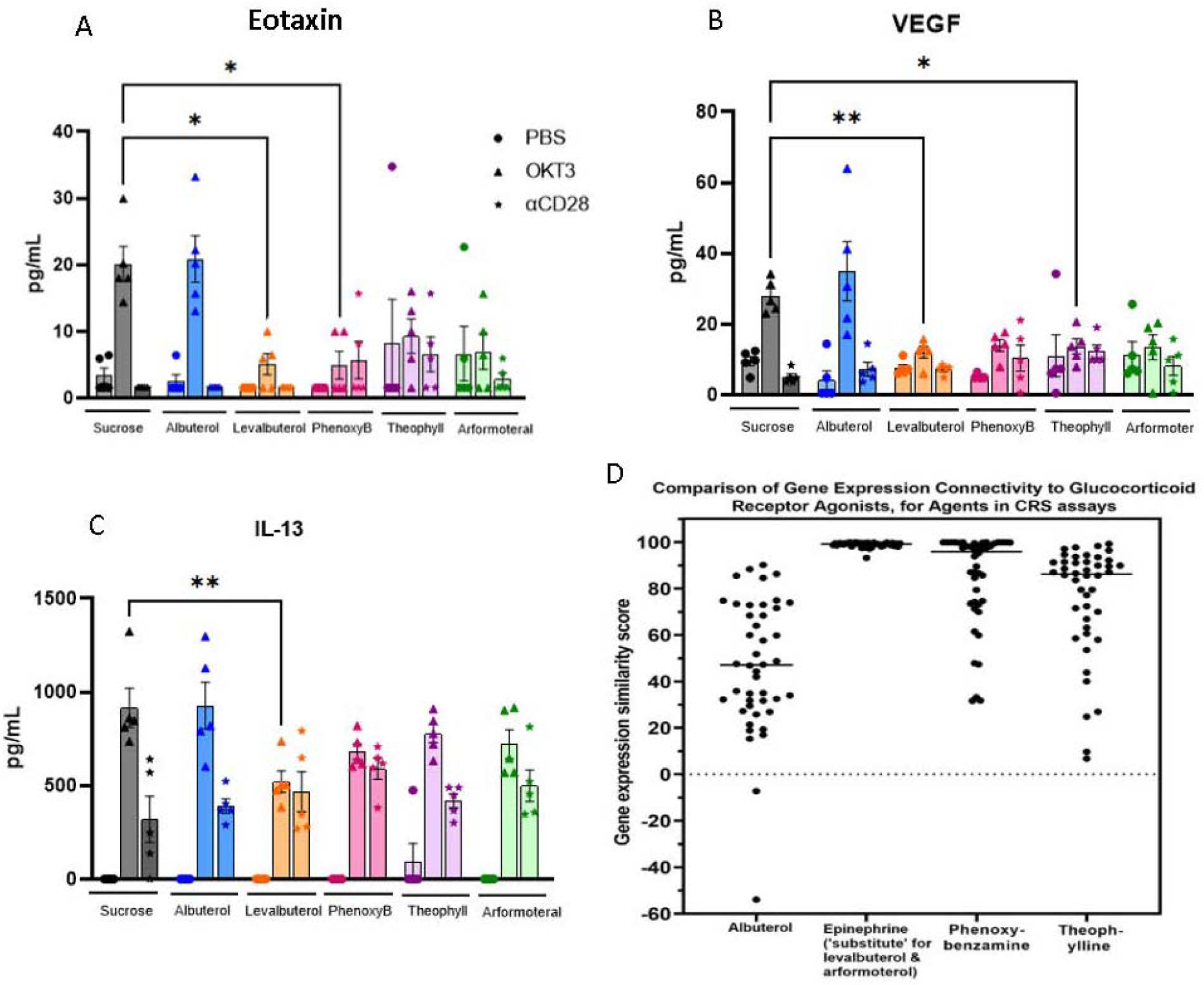
The effects of β2-adrenergic related drugs on the release of eotaxin and VEGF. Elaboration of Eotaxin (A), VEGF (B), and IL-13 (C), in the CRS assay. D, comparisons are illustrated in the same order for the gene-expression connectivity of the agents as glucocorticoid receptor agonists in the CLUE database. As noted, the gene-expression connectivity for epinephrine, the prototypical beta2-adrenergic agonist, is substituted for levalbuterol and arfomoterol since these drugs were never profiled in the CLUE database. Albuterol, the racemic mixture of R and S levalbuterol, showed no inhibition of cytokine/chemokine release in any of the assays (A, B, C).

The changes seen in cytokine and chemokine inhibition at 6 hours after OKT3 induction did not persist until assays on Day 9, 72 hours after induction. A possible explanation may be related to the fact that water intake containing the adrenergic-related pretreatments was reduced by approximately 55 to 70 percent among the treatments by Day 9 (Figure 8).

The results for eotaxin and VEGF with levalbuterol (Fig. 9) showed that some of the other beta2-adrenergic related drugs were quite close to statistical differences from their controls, so a simple “power analysis” was done by duplicating the data to double the sample size to 10 mice/group; the variance was kept the same. All the drug treatments except albuterol showed significantly less elaboration of VEGF and eotaxin with these increases in sample size (Figure 10).

**Fig 10.**
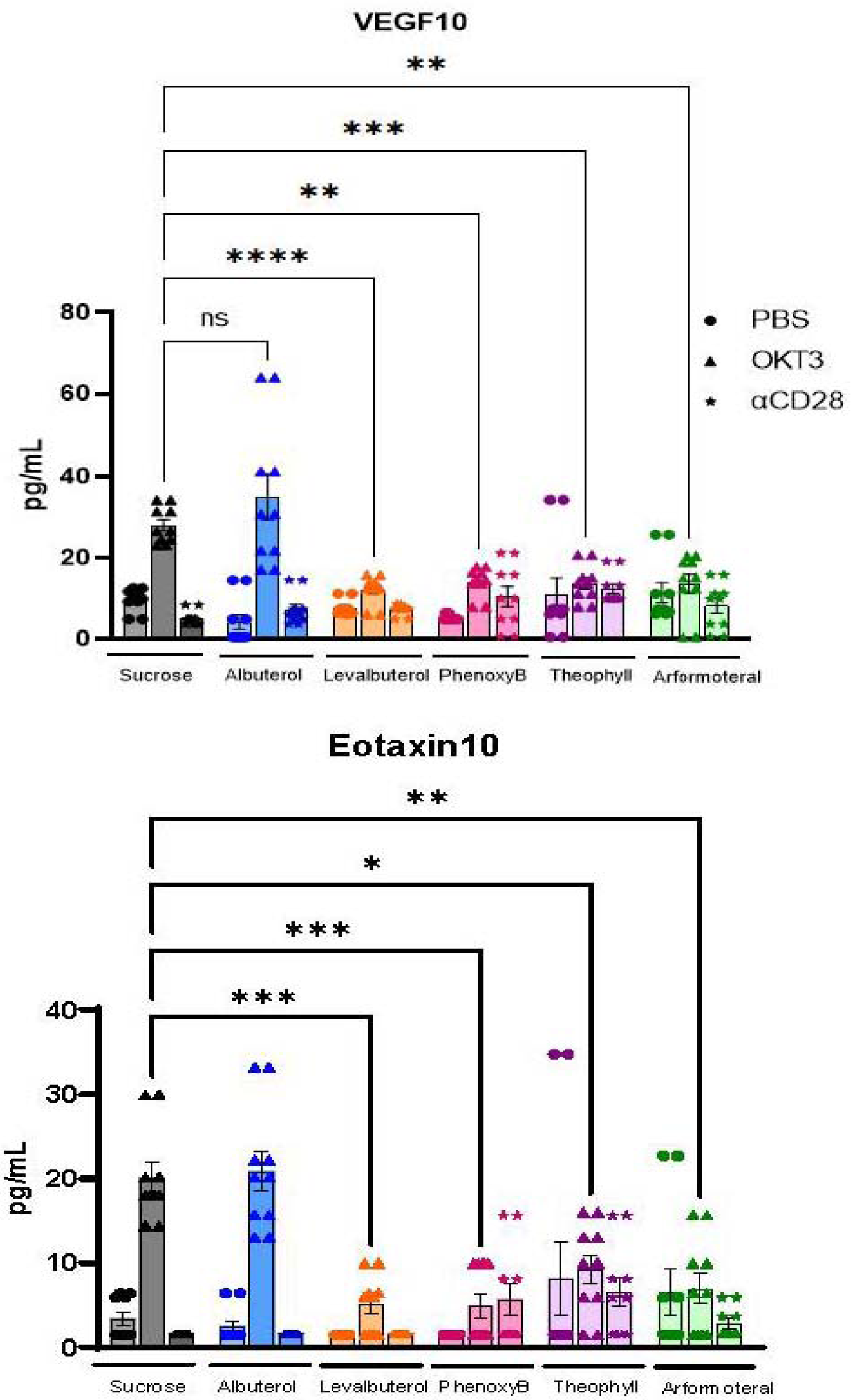
“Power” analysis of the results for the VEGF and eotaxin assays. Duplication of data from 5-animal CRS assays but variance unchanged.

Possible trends toward statistical significance were examined for all the cytokines and chemokines that were measured. Unpaired t-tests, without correction for multiple comparisons, identified the following inhibitory effects for levalbuterol on cytokine elaboration after OKT3 induction: TNFα, p = 0.0321; TNFβ, p = 0.0373; M-CSF1, p = 0.0342; IL-1α, 0.0399; and IL-18, p = 0.0003.

The MILLIPLEX panel used to quantify cytokines in the above assays identified eotaxin-1 (CCL11), which is the most extensively characterized and studied of the eotaxins. When eotaxin links to its most specific receptor, CCR3, in organs and the brain it recruits eosinophils that upon degranulation release eosinophil granule proteins, growth factors and cytokines causing cellular damage; eotaxin plays a central role in mediating a number of eosinophilic disorders, including atopic dermatitis, eosinophilia in asthma, chronic rhinosinusitis and eosinophilic esophagitis [58, 59]. Teixeira et al. [60] have reviewed the association of increased blood levels of eotaxin-1/CCL11 in patients with major psychiatric disorders including schizophrenia, major depression, bipolar disease, autism spectrum disorder, dysthymia, and Alzheimer disease; they also discussed the presence of the CCR3 receptor on microglia, and Peterson et al. [11] have proposed a pro-inflammatory role of activated microglia in destruction of dopaminergic neurons in PD. Chandra et al. demonstrated that eotaxin is markedly elevated in the substantia nigra pars compacta in post-mortem brains of PD patients compared to non-affected controls [61].

The finding that IL-13 release was also inhibited by levalbuterol (Figure 9) is of interest because IL-13 is one of the cytokines that is central to cellular trafficking; blocking its ligands is a feature of dupilumab in the treatment of chronic obstructive pulmonary disease (COPD) and other eosinophilic disorders [59, 62].

Figure 9 also presents the results for the inhibition of the release of VEGF by levalbuterol and theophylline. There is a vast literature on the importance of VEGF in neoplastic disease including an earlier comprehensive review by Goel and Mercurio [63], and most recent efforts to block certain particularly aggressive cancers that depend on binding of VEGF to a co-factor, neuropillin-2 [64]. The MILLIPLEX panel that was used to quantify this cytokine in the CRS assays was specific for the VEGF_165_ variant. Although it is many decades since the recognition of the involvement of VEGF in tumor angiogenesis, treatment complexity persists. Mabeta and Steenkamp [65] have presented a recent review of VEGF variants and VEGF_165_ is the prototypical cytokine and is upregulated in approximately 70% of tumors including breast cancer, glioblastoma multiforme, ovarian cancer, melanoma, esophageal cancer, gastrointestinal tract cancers, lung cancer, renal carcinomas and others. VEGF_165_ promotes sprouting of existing vessels as well as splitting of vessels as part of its angiogenic effects; these processes may be part of an effort by tumors to escape therapeutic interventions that produce conditions jeopardizing tumor expansion [65]. It is interesting to speculate that inhibition of release of VEGF, as seen in Figure 9, may be superior to the complexities of control of angiogenesis at the tumor level.

### 3.3. Relevant observations

The marked similarity between gene-expression connectivity for epinephrine (and presumably other β2AR agonists) and glucocorticoids, as presented in these investigations, raises the question as to why β2AR agonists have shown absolutely no evidence of the classical adverse effects of glucocorticoids. This is true with chronic treatment of β2AR agonists at pharmacological doses in their primary role as bronchodilators. Adverse side effects of glucocorticoids include hypertension, hyperglycemia, osteoporosis, glaucoma, cataract formation, peptic ulcer, gastrointestinal bleeding, and others [66]. These effects are dose dependent and reduction in dosages leads to loss of therapeutic effectiveness.

Dose limitations represent a major disadvantage in the clinical use of glucocorticoids. In addition to their efficacy as anti-inflammatory/immunomodulatory agents, these drugs have major anti-neoplastic effects in the treatment of several hematopoietic malignancies with lymphatic lineage, including chronic lymphocytic leukemia, acute lymphoblastic leukemia, multiple myeloma, Hodgkin’s lymphoma, and non-Hodgkin’s lymphoma [67, 68]. Induction of apoptosis appears to be a primary mechanism in the treatment of these malignancies [69].

There has been considerable effort directed in attempts to separate the desired effects of glucocorticoids from the initiation of adverse events. Agents of this type are non-steroidal chemically and have been variously termed as “dissociated glucocorticoid receptor ligands” [70], “selective glucocorticoid receptor agonists” (SEGRAs) [71], or examples of “biased signaling” [72]. Kleiman and Tuckerman have summarized the results with some of the biological trials with SEGRAs and they have shown limited success [73].

The fact that epinephrine, the prime example of a β2AR agonist, showed exactly the same gene-expression profile to hydrocortisone in the CLUE analyses (Tables 6 and 10; Fig. 4) but is not associated with any of the adverse effects of glucocorticoids is now known with confidence to be related to the gene-signaling pathways from different receptor ligands. The biological result is related to which gene-signaling pathways are favored [74].

An updated comprehensive review of the glucocorticoid receptor by Nicolaides et al. [75] discusses the two major classifications of the signaling of gene expression in relation to the potential for adverse events with longer treatment and higher doses versus the pathway for therapeutic predominance. The contrasting pathways are termed “transactivation” and “transrepression.” Transactivation correlates with downstream transcription of protein sequences that mediate the toxic manifestations of glucocorticoid treatment (diabetes mellitus, osteoporosis, hypertension, etc.). Transrepression is primarily associated with therapeutic effects of glucocorticoids, i.e., their anti-inflammatory, immunomodulatory, and certain anti-neoplastic effects. The mechanisms of transactivation and transrepression are complex and are discussed in extensive detail by Lesovaya et al [76]. In view of the potential anti-inflammatory effects of β2AR-related drugs observed in the present study, it would be expected that they mediate via a transrepression effect at the GR.

The chapter by Nicolaides et al. [75] also discusses the evolution of the glucocorticoid receptor, which apparently represents an important step in the survival advantage of a branch that ultimately led to mammalian species. That step, which included specificity of GR for cortisol, took place about 450 million years ago with the first appearance of the GR in ray-finned fish and then to land vertebrates. Of interest to the present study, the evolution of the sympathetic nervous system took place in that same period of geologic time, and it is presumed that a sympathetic nervous system represented a level of functional sophistication that contributed to survival to further evolution [77].

The adrenal steroids and sympathetic nervous systems are critical parts of the integrated response to stress in humans today and this may be the reason that the CLUE platform of the Harvard/MIT Broad Institute database showed essentially the same gene-expression signatures for hydrocortisone and epinephrine, as noted above. In these integrated systems, glucocorticoids have been demonstrated to promptly increase elaboration of cAMP by several mechanisms that would be expected to inhibit the release of pro-inflammatory cytokines and chemokines: ***a***) Hydrocortisone (cortisol) directly activates adenylyl cyclase to elaborate cAMP; ***b***) it increases the density of β2ARs on cell membranes of elements of the innate immune system; ***c***) it facilitates linking of those receptors to adenylyl cyclase to increase cAMP; ***d***) it preserves responsiveness (i.e. reverses tachyphylaxis) that may develop with continuous occupancy of β2ARs [54, 55, 78–82]; ***e***) and finally, there is the added fact that, as part of the stress response, hydrocortisone selectively activates the phenylethanolamine N-methyltransferease (PNMT) enzyme in the adrenal medulla to increase the synthesis of epinephrine [75, 83, 84]. It is interesting to speculate that the selective beta2-adrenergic-related agents studied in this investigation may extend the clinically therapeutic applications of the glucocorticoid-sympathetic stress response without incurring the adverse effects of higher glucocorticoid doses.

In particularly interesting observations by Flydal et al. [85] levalbuterol was identified as a small molecular “chaperone” that stabilized the enzyme, tyrosine hydroxylase. This enzyme catalyzes the rate-limiting step in the synthesis of dopamine. It is depleted or dysfunctional in the tyrosine hydroxylase deficiency syndrome and in PD. In an effort to repurpose treatments for these deficiencies, these investigators screened 1280 recognized drugs for their abilities to prevent excessive aggregation and/or misfolding of the tyrosine hydroxylase molecule, and to reduce the feedback inhibitory effects of dopamine on the activity of the enzyme. Through active site modeling, levalbuterol was proposed to bind to the same iron moiety as dopamine at the active site. However, binding by levalbuterol did not inhibit activity but did competitively reduce the feedback inhibitory effects of dopamine. These properties of levalbuterol may have value in reducing the dyskinesias associated with the chronic treatment of PD. Nineteen drugs from the initial screening showed promise but validation studies eliminated all but 4. Of those, levalbuterol demonstrated dose-related efficacy most consistently [85].

### 3.4. Clinical translation of the findings

The drugs studied in the present work are all FDA-approved medications, which could facilitate their possible translation to therapies that have relevance to neurodegenerative and neoplastic disorders. There is considerable clinical data regarding the use of levalbuterol and other beta2-adrenergic selective agents for the treatment of bronchial asthma in children and adults. The doses that were used in the present studies to test their effects on cytokine release in the human PBMC-engrafted mice were translated from human adult doses that relieve asthmatic attacks. Heart rate changes were not monitored in the mice but there were no untoward effects noted. In a clinical comparison between levalbuterol and the racemic drug albuterol in children aged 4 to 11 years in age, levalbuterol appeared to be superior in terms of improvement in respiratory function indices [86]. Children who received the recommended dose of levalbuterol for this age group (0.31 mg by nebulized inhalation) showed no mean change in heart rate; those that required double this dose for a satisfactory treatment of an asthmatic attack had a mean increase in heart rate of approximately 7 beats per minute. It must be noted that this route of administration essentially delivers the entire dose to the lungs where there is rapid systemic absorption that impacts the heart; this would not be the case if the drug was administered episodically by the oral route. It was administered continuously in drinking water in the mouse studies. A recent meta-analysis covering the use of this class of drugs from 1996 – 2021 demonstrated their safety [87].

Clinical translation would also be aided by the fact that there are no formulation issues for levalbuterol. There are three FDA-approved dose levels that could be used for episodic administration orally.

### 3.5. Limitations

An unfortunate limitation in the studies, as noted above, was the fact that levalbuterol and arformoterol have never been profiled for gene-expression connectivity to glucocorticoids by the Harvard/MIT Broad Institute. We did formally nominate it for assay in CLUE, but this has not taken place. This remains to be addressed in future studies.

Another limitation was that we were not able to assay the proto-typical beta2-adrenergic agonist, epinephrine, in the mouse CRS assay. The assay was designed to provide a low-stress delivery of pretreatments to the mice by providing the drugs in the drinking water. Epinephrine would undergo autoxidation under these conditions.

## 4. Conclusions

Advances in the development of beta2-selective adrenergic agonists has led to their recognition as potent anti-inflammatory agents with potential for treatment in Parkinson’s disease and inflammatory disorders. This potential was reinforced in the present study by observations that epinephrine, the prototype beta2-adrenergic agonist, showed an almost identical gene expression signature as cortisol (hydrocortisone) and other glucocorticoids.

Levalbuterol and other beta2-adrenergic-related agents inhibited the release of eotaxin, VEGFa and IL-13 from mice engrafted with human peripheral blood monocyte cultures in a Jackson Laboratory assay, supporting the predictions of similar gene expression mechanisms and potential translation to inflammatory pathologies and neoplastic syndromes. The observations with the group of drugs that were studied suggested a unifying mechanism of action of maintenance or elevation in cellular levels of cAMP. This aspect requires further investigation.

While therapeutic efforts in neurodegenerative, inflammatory, and neoplastic diseases that block downstream pathways and receptors have advanced considerably, there remain the issues of development of tolerance, escape mechanisms and off-target adverse effects. There may be a philosophical advantage to inhibition of the release of inciting elements in these pathologies, as suggested by the present studies.

The agents studied are FDA-approved drugs with minimal adverse effect profiles. The fact that they showed no toxicity in the animal studies at doses translated from human experience, and that there are existing formulations that can be administered orally, may facilitate their repurposing and ultimate translation to appropriate pathologies.

## Supporting Information

**Figure S1:**
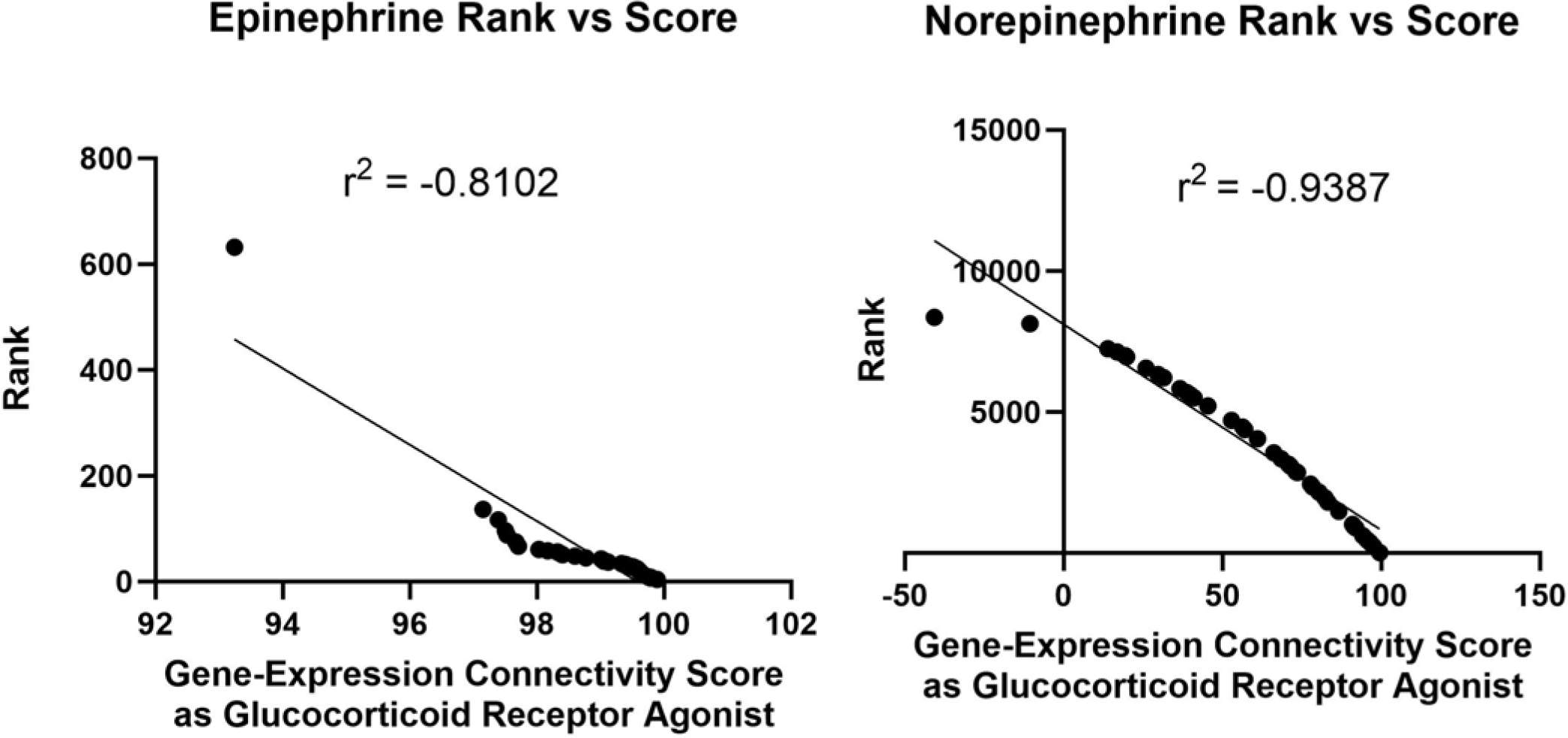
Association between rank score and gene-expression connectivity score.

Comparison of the associations between the Rank and the Gene-Expression Connectivity Scores for glucocorticoid receptor agonist activity for epinephrine and norepinephrine in the CLUE database. Both associations show highly significant inverse relationships for the two measures. However, the beta2-adrenergic prototype, epinephrine, shows a much closer relationship between rank and the gene-expression connectivity scores. Norepinephrine lacks significant beta-2 adrenergic activity (Table 8). The data for these two measures for epinephrine and norepinephrine are presented in Tables 6 and 7, respectively. A plot demonstrating these differences is presented in Figure 3. In practical terms, “rank” represents the relative depth that the CLUE database must be probed to find gene-expression connectivity with perturbagens; the further the probe, the weaker the connectivity.

**Table S1:**
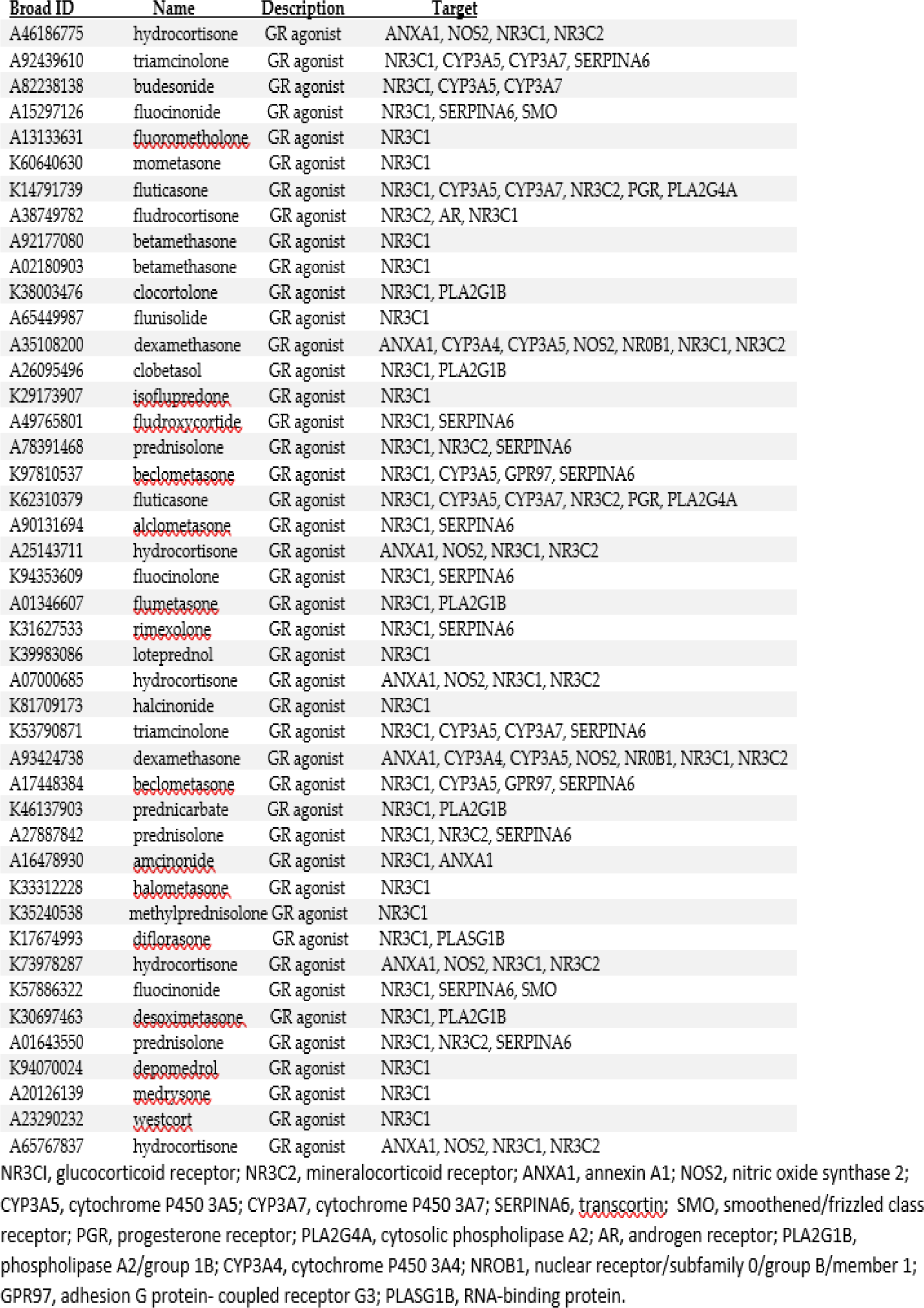
Glucocorticoid Receptor (GR) Agonist Class List with Targets.

**Table S2:**
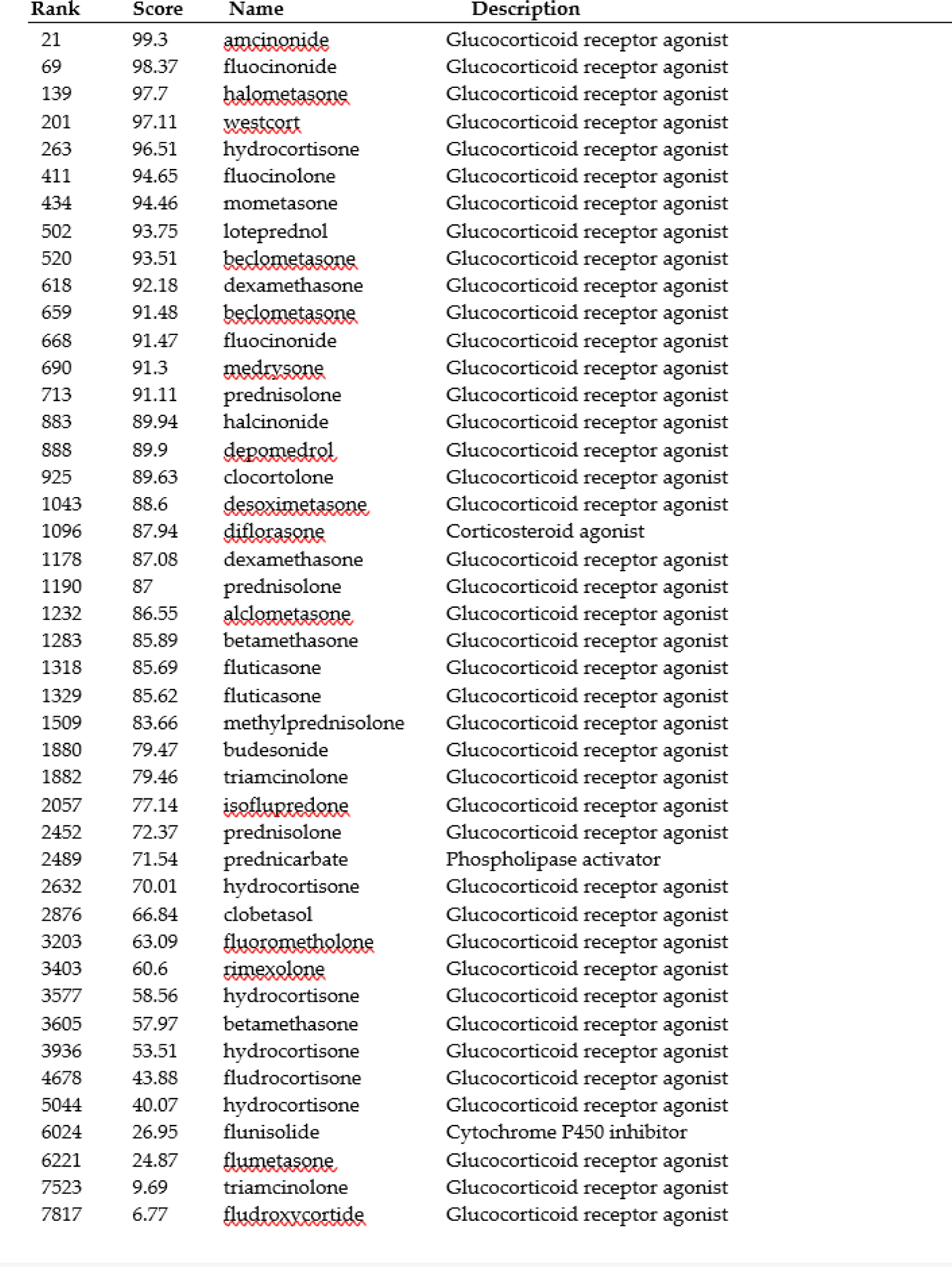
Gene expression connectivity scores for THEOPHYLLINE vs GR agonists.

**Table S3:** Gene expression connectivity scores for CAFFEINE vs GR agonists.

| Rank | Score | Name | Description |
| --- | --- | --- | --- |
| 213 | 98.34 | alclometasone | Glucocorticoid receptor agonist |
| 266 | 98.06 | beclomethasone | Glucocorticoid receptor agonist |
| 294 | 97.84 | fluticasone | Glucocorticoid receptor agonist |
| 340 | 97.74 | rimexolone | Glucocorticoid receptor agonist |
| 342 | 97.74 | budesonide | Glucocorticoid receptor agonist |
| 369 | 97.6 | diflorasone | Corticosteroid agonist |
| 546 | 96.38 | desoximetasone | Glucocorticoid receptor agonist |
| 605 | 95.96 | prednisolone | Glucocorticoid receptor agonist |
| 801 | 94.26 | isoflupredone | Glucocorticoid receptor agonist |
| 802 | 94.26 | betamethasone | Glucocorticoid receptor agonist |
| 984 | 92.8 | triamcinolone | Glucocorticoid receptor agonist |
| 1146 | 91.13 | clocortolone | Glucocorticoid receptor agonist |
| 1238 | 90.46 | loteprednol | Glucocorticoid receptor agonist |
| 1297 | 89.85 | methylprednisolone | Glucocorticoid receptor agonist |
| 1357 | 89.37 | prednicarbate | Phospholipase activator |
| 1525 | 87.34 | dexamethasone | Glucocorticoid receptor agonist |
| 1924 | 82.44 | prednisolone | Glucocorticoid receptor agonist |
| 1990 | 81.65 | dexamethasone | Glucocorticoid receptor agonist |
| 2060 | 80.88 | beclomethasone | Glucocorticoid receptor agonist |
| 2128 | 80.38 | fluocinolone | Glucocorticoid receptor agonist |
| 2164 | 79.94 | flumetasone | Glucocorticoid receptor agonist |
| 2431 | 77.42 | hydrocortisone | Glucocorticoid receptor agonist |
| 2515 | 76.32 | clobetasol | Glucocorticoid receptor agonist |
| 3163 | 67.41 | halometasone | Glucocorticoid receptor agonist |
| 3761 | 58.11 | fludroxycortide | Glucocorticoid receptor agonist |
| 3832 | 57.19 | westcort | Glucocorticoid receptor agonist |
| 3839 | 57.12 | prednisolone | Glucocorticoid receptor agonist |
| 3873 | 56.65 | medrysone | Glucocorticoid receptor agonist |
| 4311 | 50.09 | depomedrol | Glucocorticoid receptor agonist |
| 4632 | 45.21 | mometasone | Glucocorticoid receptor agonist |
| 4731 | 44.04 | fluocinonide | Glucocorticoid receptor agonist |
| 4870 | 42.07 | hydrocortisone | Glucocorticoid receptor agonist |
| 4954 | 40.88 | hydrocortisone | Glucocorticoid receptor agonist |
| 5046 | 39.65 | hydrocortisone | Glucocorticoid receptor agonist |
| 5467 | 34.15 | amcinonide | Glucocorticoid receptor agonist |
| 5826 | 28.63 | fluorometholone | Glucocorticoid receptor agonist |
| 5924 | 27.31 | triamcinolone | Glucocorticoid receptor agonist |
| 6059 | 24.86 | halcinonide | Glucocorticoid receptor agonist |
| 6203 | 22.66 | betamethasone | Glucocorticoid receptor agonist |
| 6372 | 19.84 | flunisolide | Cytochrome P450 inhibitor |
| 6673 | 15.86 | hydrocortisone | Glucocorticoid receptor agonist |
| 6678 | 15.81 | fluticasone | Glucocorticoid receptor agonist |
| 7332 | 5.38 | fludrocortisone | Glucocorticoid receptor agonist |
| 7492 | 3 | fluocinonide | Glucocorticoid receptor agonist |

### Striking Image

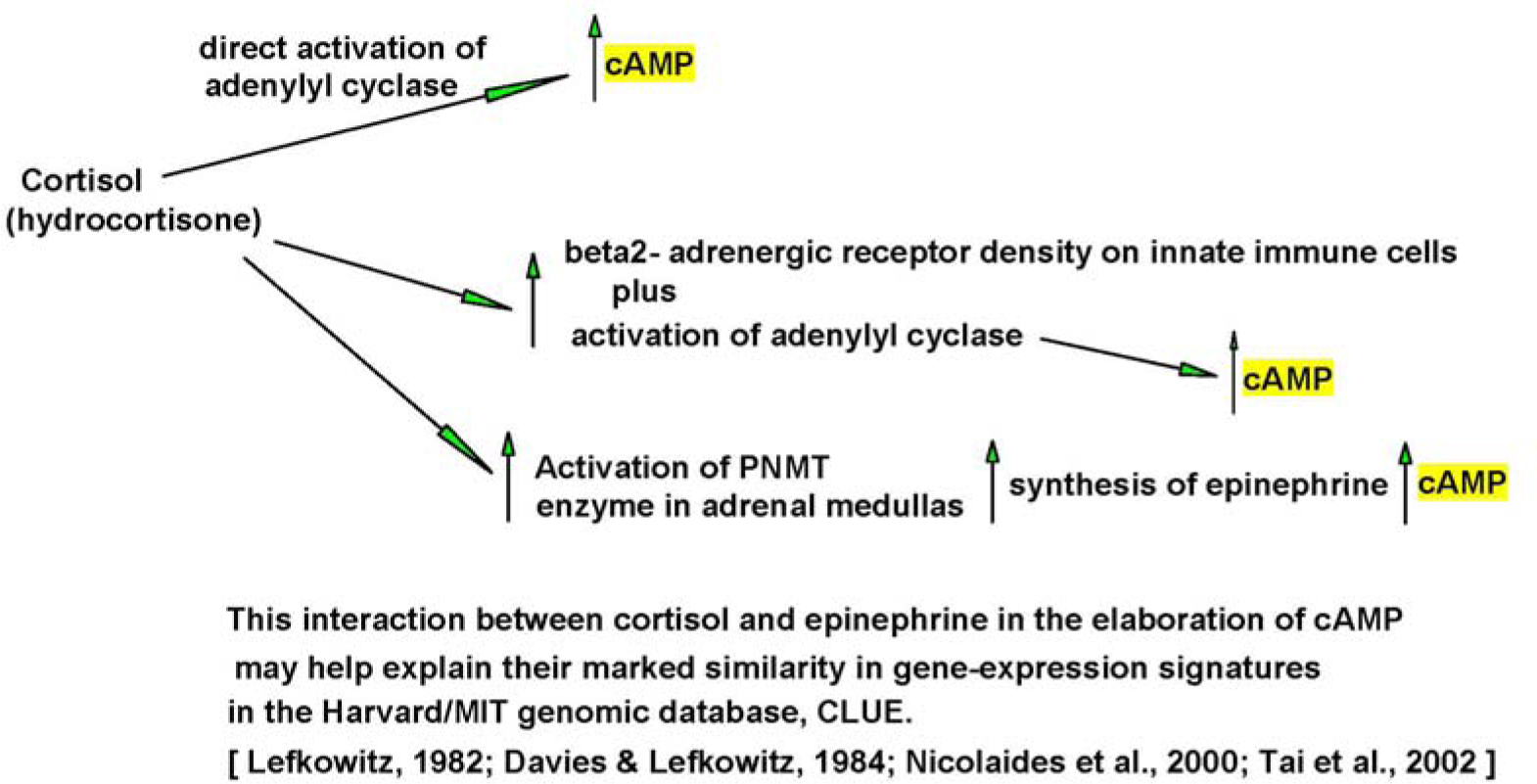

## Author Contributions

The author conceived the investigation, carried out bioinformatics studies, contributed to the design of the animal assays, and wrote the manuscript.

## Data availability

Essentially all the data are included in the manuscript. Body weights for individual animals and water consumption data by cage are available upon request.

## Acknowledgments

The author is grateful to Ms. Jennifer Brown for editorial assistance in the preparation of the manuscript, and to Ms. Brown and Ms. Gail Anderson of the Department of Pharmacology, New York Medical College, for facilitating the administrative and financial procedures required for the studies. The author also recognizes the contributions of Ms. Maribeth Balingit and associates of the Phillip Capozzi, M.D. Library, New York Medical College, for assistance in assembling the bibliography of the manuscript.

